# Deciphering the regulatory logic of a *Drosophila* enhancer through systematic sequence mutagenesis and quantitative image analysis

**DOI:** 10.1101/2020.06.24.169748

**Authors:** Yann Le Poul, Yaqun Xin, Liucong Ling, Bettina Mühling, Rita Jaenichen, David Hörl, David Bunk, Hartmann Harz, Heinrich Leonhardt, Yingfei Wang, Elena Osipova, Mariam Museridze, Deepak Dharmadhikari, Eamonn Murphy, Remo Rohs, Stephan Preibisch, Benjamin Prud’homme, Nicolas Gompel

**Affiliations:** Chair of Evolutionary Ecology, Ludwig-Maximilians Universität München, Fakultät für Biologie, Biozentrum, Grosshaderner Strasse 2, 82152 Planegg-Martinsried, Germany; Berlin Institute for Medical Systems Biology, Max Delbrück Center for Molecular Medicine, Robert-Rössle-Str. 10, 13092 Berlin, Germany; Chair of Human Biology and Bioimaging, Ludwig-Maximilians Universität München, Fakultät für Biologie, Biozentrum, Grosshaderner Strasse 2, 82152 Planegg-Martinsried, Germany; Aix-Marseille Université, CNRS, IBDM, Institut de Biologie du Développement de Marseille, Campus de Luminy Case 907, 13288 Marseille Cedex 9, France; Quantitative and Computational Biology, Departments of Biological Sciences, Chemistry, Physics & Astronomy, and Computer Science, University of Southern California, Los Angeles, California 90089, USA; Janelia Research Campus, Howard Hughes Medical Institute, VA, Ashburn, USA

## Abstract

Transcriptional enhancers are short DNA sequences controlling the spatial activity, timing and levels of eukaryotic gene transcription. Their quantitative transcriptional output is thought to result from the number and organization of transcription factor binding sites (TFBSs). Yet, how the various aspects of regulatory information are encoded in enhancer sequences remains elusive. We addressed this question by quantifying the spatial activity of the *yellow spot* enhancer active in developing *Drosophila* wings. To identify which enhancer DNA sequence contributes to enhancer activity, we introduced systematic mutations along the enhancer. We developed an analytic framework that uses comprehensive descriptors to quantify reporter assay in transgenic flies and measure spatial variations in activity levels across the wing. Our analysis highlights an unexpected density of regulatory information in the *spot* enhancer sequence. Furthermore, it reveals an unanticipated regulatory logic underlying the activity of this enhancer, and how it reads the wing *trans*-regulatory landscape to encode a spatial pattern.

## Introduction

Enhancers constitute a particular class of *cis*-regulatory elements that regulate in which cells a gene is transcribed, when, and at which rate (Banerji et al., 1983; Levine, 2010; Shlyueva et al., 2014). Notably, enhancers play a central role during development in plants and animals (Peter and Davidson, 2015), integrating spatial information from transcription factors (TFs) bound to them. The resulting transcriptional activity provides a blueprint for morphology. In spite of its central role in the construction of multicellular organism, the regulatory logic underlying an enhancer’ spatial activity remains largely elusive. The transcription machinery associated with a core promoter samples TFs bound to a distant enhancer through the mediator complex and in turn initiates transcription (Levine, 2010). Hence, the mode of TF sampling represents an important step for enhancer activity and is critically linked to the organization of TF binding sites (TFBSs) in the enhancer sequence (Weingarten-Gabbay and Segal, 2014). Deciphering the regulatory logic of an enhancer, therefore, means understanding the causal relationship between the composition of its DNA sequence and the resulting spatio-temporal transcriptional activity from its target promoter. Understanding the regulatory logic means, in particular, to be able to design custom sequences capable of specific transcriptional activities, a goal that is hardly reached (Crocker et al., 2017; Kulkarni and Arnosti, 2003; Ronchi et al., 1993; Vincent et al., 2016).

In terms of sequence composition, the main focus is on the organization of TFBSs: their nature (activating, repressing, granting accessibility), their number, their orientation and spacing, their relative affinity for particular TFs, as well as the interactions of their cognate TFs (cooperative binding, synergy, competition, short-and long-range repression) (Barolo, 2016; Erceg et al., 2014; Long et al., 2016; Weingarten-Gabbay and Segal, 2014; Yanez-Cuna et al., 2013). TFBS organization has been under intense scrutiny in the last two decades (Evans et al., 2012; Hare et al., 2008; Khoueiry et al., 2010; Levo and Segal, 2014; Ludwig et al., 2000; Ludwig et al., 2011; Spitz and Furlong, 2012; Swanson et al., 2011). Two opposing models, the enhanceosome model, which posits a highly constrained organization of TFBSs to favor extensive interactions between bound TFs, and the billboard model, which assumes a flexible organization because bound TFs function in a largely independent manner, have dominated the literature on enhancers and have been extensively discussed elsewhere (Arnosti and Kulkarni, 2005; Kulkarni and Arnosti, 2003; Ludwig et al., 2011; Ludwig et al., 2005; Panne et al., 2007; Slattery et al., 2014; Spitz and Furlong, 2012; Swanson et al., 2011; Thanos and Maniatis, 1995). Both models are based on the transcriptional consequences resulting from perturbations to the TFBS organization. Implicitly, they focus on known TFBSs and pay little attention to the remaining sequences. In most cases, however, available TFBS information has been insufficient to create functional synthetic enhancers (Vincent et al., 2016). It appears, therefore, that uncovering the determinants of enhancer activity requires, along with a classical mutational approach, a systematic testing of sufficiency.

In terms of transcriptional readout, the assessment of an enhancer’s activity is often limited to describing qualitative spatial patterns. This is in part because the sole qualitative spatial expression of numerous patterning genes during development is seen as a surrogate for the final morphology (Carroll, 1998). Yet, the overall transcriptional levels of developmental genes must be tightly controlled for normal development, as gain-of-function and hypomorph mutants, or RNA-seq experiments show. A number of studies have taken activity levels into account in individual cells in cell culture experiments (*e*.*g*., (King et al., 2020; Kircher et al., 2019; Kwasnieski et al., 2012)) or dissociated tissue (*e. g*., (Farley et al., 2015)), but in this case the information on spatio-temporal variation (the pattern), is lost. By contrast, other studies have quantified pattern elements of enhancer activity but with limited spatial resolution (Crocker et al., 2015; Crocker and Stern, 2017; Dufourt et al., 2018)). It is nevertheless important to appreciate that the overall levels and the spatial pattern of activity in a given tissue are intrinsically linked. To understand how the information of TFs is integrated in an enhancer to produce an activity, it is therefore critical to measure simultaneously and comprehensively variations in both activity pattern and levels with quantitative descriptors.

In this work, we present an agnostic approach to decipher the logic of a regulatory element of the gene *yellow* from the fruit fly *Drosophila biarmipes*, the so-called *spot*^*196*^ enhancer, driving patterned gene expression in pupal wings (Arnoult et al., 2013; Gompel et al., 2005). We introduced systematic small-scale mutations of different sizes along the 196 base pairs (bp) of the enhancer sequence to test the necessity of the mutated positions, and large block randomization to test sufficiency of the intact sequence to drive an activity, without *a priori* functional assumptions on the mutated nucleotides, to uncover the regulatory logic underlying the activity of the *spot*^*196*^. To assess the activity of each mutant enhancer, we devised a pipeline that uses comprehensive descriptors to quantify reporter assay in transgenic flies and measure variations in activity level across the fly wing (Le Poul et al., in preparation). Combining enhancer mutational dissection at different scales with quantitative spatial activity analyses, we evaluated how altering the enhancer structure affects the activity across the regulatory landscape of the wing, and the sufficiency of regulatory units to drive the different components of the activity.

Our results show that, surprisingly, all mutated positions along the sequence contribute significantly to the regulatory logic. Multiple previously unknown sites showed minor to major effects, including successions of repressing and activating segments, unraveling a high density of regulatory information. The sufficiency analysis also uncovered the primary determinant of activity in segments of moderate effects. Altogether, we reveal an unanticipated regulatory logic, where the simple spatial pattern of enhancer activity in the wing results from a complex interplay between activators, repressors, and repressors of repressors.

## Results

### A quantitative framework to dissect a regulatory element active in *Drosophila* wings

To assess how the sequence of an enhancer relates to its regulatory activity, we used a recently evolved enhancer in *Drosophila*. Several fruitfly species have gained patterns of dark pigmentation through a change in developmental expression of pigmentation genes such as *yellow* during metamorphosis (Wittkopp et al., 2003). For instance, the gain of a spot of dark pigmentation on the wings of *Drosophila biarmipes* entails the evolution of a new regulatory element, the *spot* enhancer at the *yellow* locus, upstream of the transcription start site (Gompel et al., 2005). The dissection of this enhancer has shown that a minimal 196 bp fragment, *spot*^*196*^, was sufficient to drive reporter expression in a proximal-distal gradient toward the distal wing tip in transgenic *D. melanogaster* (Gompel et al., 2005). It contains at least 4 TFBSs for the activator Distal-less (Dll) and at least one TFBS for the repressor Engrailed (En) (Arnoult et al., 2013; Gompel et al., 2005) (Figure 1A). Together, these inputs were considered to be sufficient to explain the spatial activity of *spot*^*196*^ in the wing, with activation in the distal region and repression in the posterior wing compartment (Arnoult et al., 2013; Gompel et al., 2005). Grafting TFBSs for these factors on a naive sequence in their native configuration, however, proved insufficient to produce regulatory activity in wings (B. Prud’homme and N. Gompel unpublished results). This prompted us to dissect the *spot*^*196*^ element further, to identify what determines its regulatory activity. To this end, we moved away from qualitative appraisal of mutant enhancer activity (Arnoult et al., 2013; Gompel et al., 2005) and devised a quantitative assay and analytic tools to examine the activity of mutants of this enhancer in reporter constructs in transgenic *D. melanogaster* (here used as an experimental recipient with site-specific integration). First, to fully and precisely appreciate the effect of mutations on the spatial and quantitative activity of the enhancer, we have developed a technical and conceptual framework for measuring reporter expression levels and comparing them among constructs (see methods for details). In brief, for each reporter construct line, we sampled over 30 male wings (one wing per fly). We imaged each wing with a wide-field microscope under bright-field and fluorescent light, and detected the venation on the bright-field images of all wings. To compare specimens, we then assessed reporter activity relative to the venation structure detected on every wing. To this end, we used a deformable model (thin plate spline) to warp the fluorescent image of each subject, using landmarks placed along the veins of the corresponding bright-field image, and aligning them to a reference venation. The resulting dataset is a collection of fluorescence images for which the venation of all specimens is perfectly aligned. These images, represented as the list of fluorescence intensity of all pixel, constitute the basis of all our quantitative dissection. To assess whether or not the activity driven by a given mutant significantly differs from that of the wild type, or any other mutant, we used the scores produced by Principal Component Analysis (PCA) that comprehensively summarizes the variation of the pixel intensities across wings. To visualize the reporter activity per line, we used images representing the average activity per pixel (hereafter: average phenotype; *e*.*g*., Figure 1B). Differential reporter gene expression is generally represented using log ratios (Robinson et al., 2010), which measure the fold changes in expression level of a gene relative to a reference (*e*.*g*., the expression of the same gene under different conditions). We applied this principle to visually compare, across the wing, differential activities between constructs by computing the log ratio between their average phenotypes at every pixel (hereafter noted *logRatio*). *logRatio* images of mutants *vs*. wild type are of particular interest to decipher the regulatory logic, because they reveal in which proportion a mutant affects the enhancer activity across the wing. It therefore represents, independently of the wild-type pattern, the effect of a mutation on the integration of spatial information operated by the enhancer. In the ideal case where a particular TF input is integrated directly and linearly with respect to the TF concentration, and the action of this TF is only modulated by uniformly distributed TFs, then the *logRatio* image reflects the spatial distribution of this particular TF. The underlying logic is straightforward: in this ideal case, a sequence mutation breaking the interaction between the DNA and the TF will have a significant effect on the phenotype. The intensity of the local phenotypic effect (relatively to the wild-type levels) will depend on the local intensity of the TF-DNA interaction across the wing: where this interaction is not happening, no effect on the phenotype is expected, and reciprocally. For any situation departing from these ideal conditions, the resemblance between the *logRatio* and the TF distribution is compromised. For instance, when a TF is locally repressed by another, *logRatio* will correspond to the net loss of spatial information integration, including the loss of TF-TF interactions. The *logRatio* of a mutant affecting a known TFBS for which the corresponding TF distribution is known therefore informs us on its contribution in the regulatory logic of the enhancer, and how linearly this integration happens. Hence, even without additional knowledge on the regulatory logic and TF spatial variation, the variety of *logRatio* patterns informs us on the minimal number of spatial inputs integrated by the enhancer, while also reflecting the valence and the magnitude of their effects.

**Figure 1.**
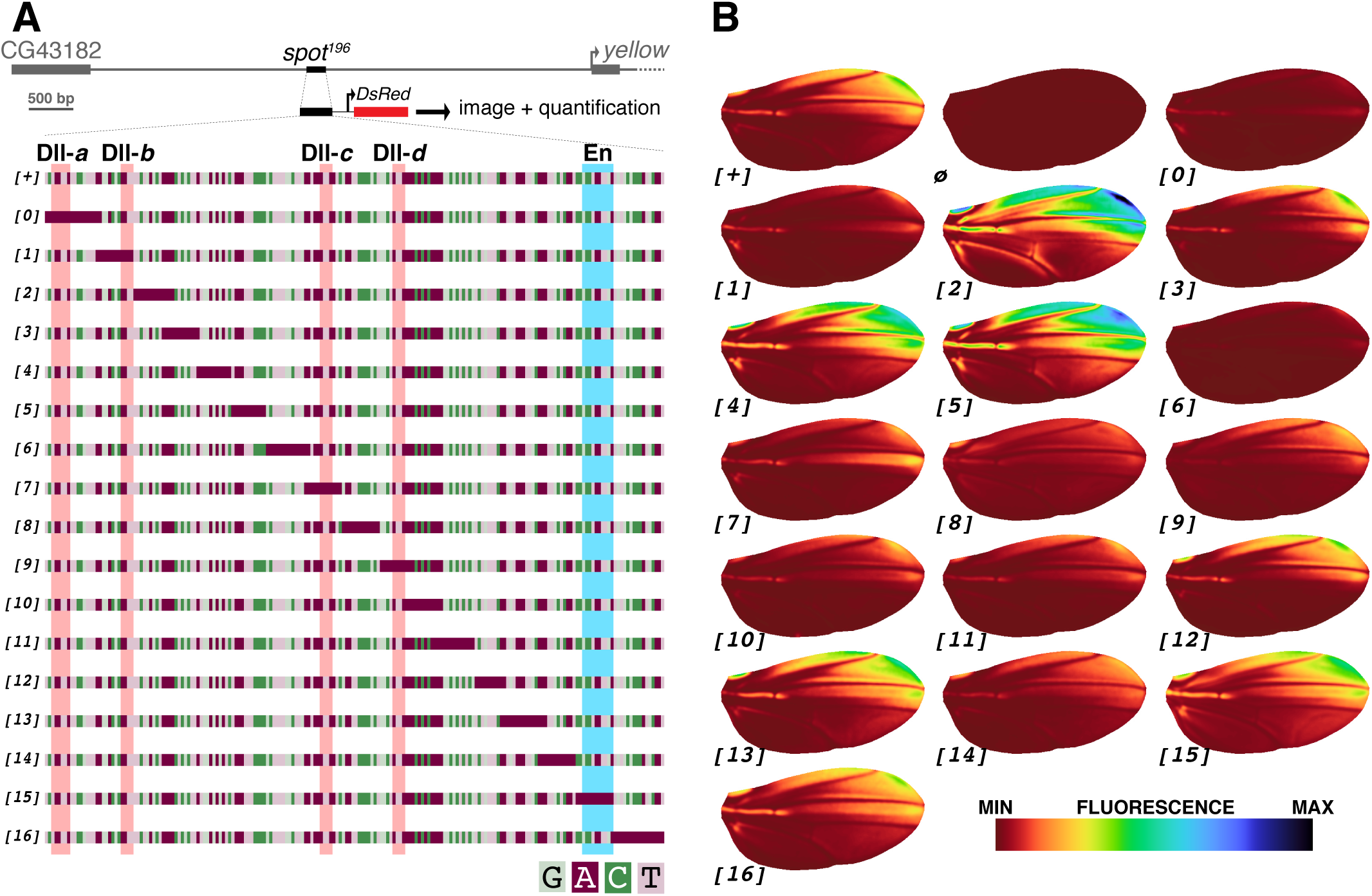
A mutational scan of the *Drosophila biarmipes spot*^*196*^ enhancer with a quantitative reporter assay. (**A**) Wild-type (*[+]*) and mutant (*[0]* to *[16]*) versions of the *spot*^*196*^ enhancer from the *D. biarmipes yellow* locus (depicted at the panel top) were cloned upstream of a DsRed reporter to assay their respective activities in transgenic *D. melanogaster*. Each mutant targets a position of the enhancer where the native sequence was replaced by an A-tract (color-code: light green=guanine, purple=adenine, dark green=cytosine, pink=thymine). Four characterized binding sites for the TF Distal-less (Dll-a, Dll-b, Dll-c and Dll-d) (Arnoult et al., 2013) are highlighted in red and a single binding site for the TF Engrailed (Gompel et al., 2005) is highlighted in blue across all constructs. (**B**) Average wing reporter expression for each construct depicted in (**A**) and an empty reporter vector (ø). Each wing image is produced from 11 to 77 individual wing images (38 on average; Figure 1–figure supplement 4), aligned onto a unique wing model. The average image is smoothened and intensity levels are indicated by a colormap.

### Every nucleotide position along the *spot*^*196*^ enhancer contributes to its quantitative spatial activity in the wing

To determine the necessity of regulatory integration along the *spot*^*196*^ enhancer sequence, we generated a first series of mutants scanning the element. We maximized the disruption of sequence information by introducing stretches of 10-18 bp (11.5 bp on average) of poly(dA:dT), also known as A-tracts (Neidle, 2010) at adjacent positions along the sequence (Figure 1A). Thus, the sequence of each of the 17 constructs (*spot*^*196 [0]*^ to *spot*^*196 [16]*^, or *[0]* to *[16]* in short, Figure 1A) is identical to the wild-type *spot*^*196*^ (*[+]* in short), except for one segment where the sequence was replaced by the corresponding number of adenines. These mutations affect the local sequence composition, without changing distances or helical phasing in the rest of the enhancer. We measured activities of the respective mutant enhancers in the wing, as described above. The activity of each mutant (presented in Figure 1B) differs significantly from that of *[+]*, as measured in the PCA space (Figure 1–figure supplement 2 and 3). This means that the activity of each mutant had some features, more or less pronounced, that significantly differentiates its activity from *[+]*, revealing the high density of regulatory information distributed along the sequence of *spot*^*196*^. The magnitude and direction of the effects, however, varies widely among mutants, ranging from activity levels well above those of *[+]* to a near complete loss of activity.

The average activity levels of each mutant construct in the wing relative to the average activity levels of *[+]* show how effect directions and intensities are distributed along the enhancer sequence (Figure 2). This distribution of regulatory information, the magnitude and the direction of the effects, including several successions of over-expressing and under-expressing mutants, suggest a more complex enhancer structure than previously thought (Gompel et al., 2005).

**Figure 2.**
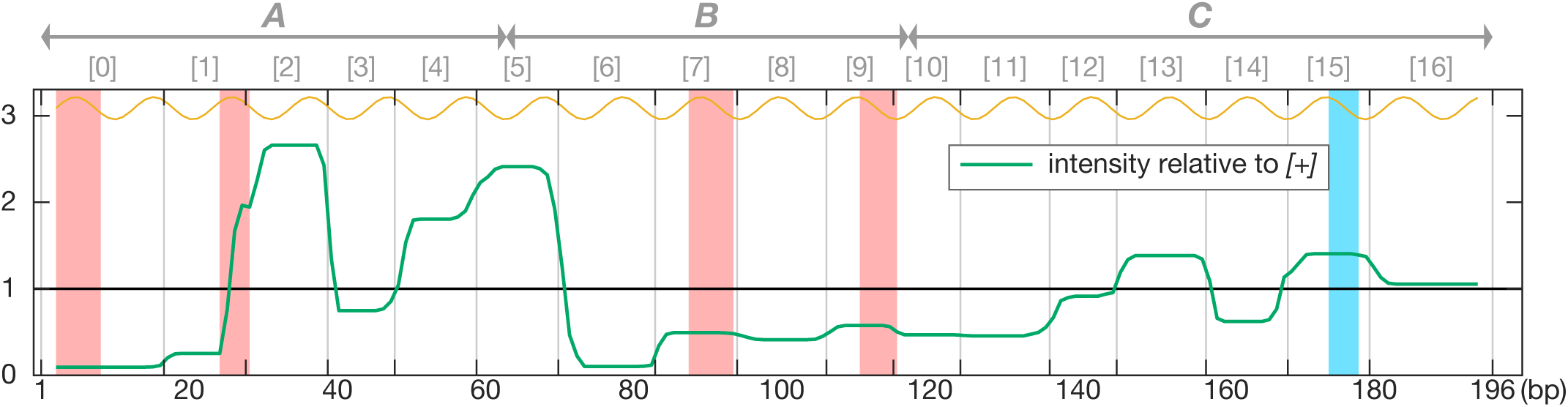
Mutational effect on intensity of activity along the *spot*^*196*^ sequence. The phenotypic effect of each mutation described in Figure 1 along the *spot*^*196*^ sequence (x-axis) is plotted as the average intensity difference to the wild type. 1 on the y-axis represents the mean wild-type intensity of reporter expression. The graph shows how each construct departs from the wild-type activity (see methods). Mutation positions in constructs *[0]*-*[16]* are indicated above the graph. The locations of blocs *A, B* and *C*, analyzed in figure 5 are also indicated above the graph. The yellow curve above the graph indicates the helical phasing.

In principle, the localized mutations we introduced can affect the *spot*^*196*^ enhancer function through non-exclusive molecular mechanisms. First, a mutation may affect TF-DNA interactions in different ways. It can directly disrupt one or more TFBS cores. For example, segments *[0], [1], [7]* and *[9]* remove characterized TFBSs for Dll and produce a reduced activity in the spot region, consistently with previous work (Arnoult et al., 2013). Segment *[15]* also suppresses a characterized TFBS for a different TF, En, and results in higher activity in the posterior wing compartment, also consistent with previous work (Gompel et al., 2005) (Figure 1B). Likewise, mutations can influence TF binding at neighboring TFBSs, as TF affinity can be affected by the nature of sequences around the TFBS (Gordan et al., 2013; Slattery et al., 2014; Yella et al., 2018), particularly by altering DNA shape properties (Abe et al., 2015; Hizver et al., 2001; Yella et al., 2018). A-tract mutations may also influence nucleosome occupancy and positioning and thereby the binding of TFs at adjacent sites (Barozzi et al., 2014). Second, and not exclusively, because they result in a local increase of DNA rigidity (Neidle, 2010; Nelson et al., 1987; Suter et al., 2000), A-tract mutations may hinder or modulate TF interactions. Such changes in rigidity, which we have evaluated for our mutant series (Fig. 3A), may affect TF-TF interactions (Fig. 3B) such as close proximity cooperative binding (Morgunova and Taipale, 2017), cooperative recruitment of a cofactor (Lim et al., 2003), or short-range repression (Gray and Levine, 1996). To evaluate the plausibility of these different possible effects, we examined more carefully the phenotypes of each mutant construct.

**Figure 3.**
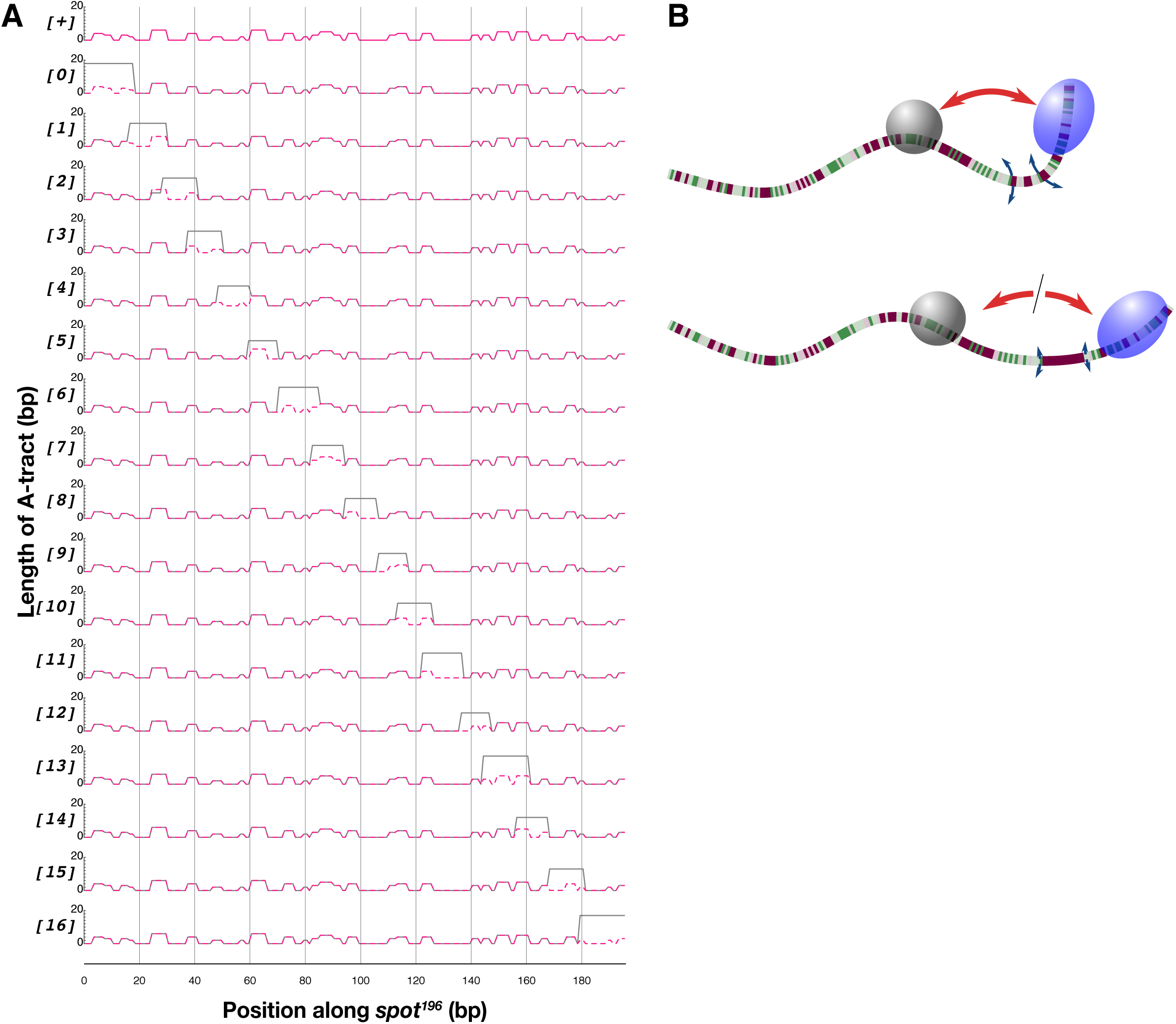
Local rigidity along the wild-type and mutant *spot*^*196*^. (**A**) Each graph is a plot of the length of the longest consecutive A_n_T_n_ sequence that a base pair participates in, a proxy for sequence rigidity at this position. The first graph on top is the wild type (*[+]*) alone. The remaining graphs show plots for each mutant (*[0]*, …, *[16]*) with a solid black line, compared to the wild type represented with a dotted magenta line. (**B**) Schematics illustrating the hypothetical consequence of local DNA rigidity (caused by an A-tract) on TF interactions. A flexible linker between two TFBSs would favor interactions between 2 bound TFs, while a stiffer linker of the same length would limit, or prevent these interactions.

### An enhancer’s view on the wing *trans*-regulatory landscape revealed by *logRatio* images

To decipher the regulatory logic of *spot*^*196*^, we first precisely analyzed the effect of mutants individually. The enhancer directly or indirectly integrates information from factors upstream in the regulatory network. Factors imparting this information are likely to have different expression patterns. Each location on the wing therefore represents a specific *trans*-regulatory environment that the enhancer integrates to produce its activity output at this location. With an average of 11.5 bp, our A-tract mutations are somewhat larger than an average eukaryotic TFBS (∼10 bp (Stewart et al., 2012)) and each mutation is likely to affect up to two TFBSs. This size represents the limit of regulatory content that we can discriminate in this study. To visualize the changes in spatial activity caused by each mutation, we compared mutants to *[+]* with *logRatios. logRatio* images represent how mutating a stretch of sequence affects the enhancer view of the wing *trans*-regulatory landscape, and can therefore, in the ideal case discussed above, reflect the distribution of the individual spatial inputs received and integrated along the *spot*^*196*^ sequence.

*logRatios* images can be particularly informative when both a TFBS and the distribution of the cognate TF are known, as it informs on how directly the TF information is integrated. This is the case for En and Dll, for which TFBSs have been previously characterized in the *spot*^*196*^ (Arnoult et al., 2013; Gompel et al., 2005). The disruption of an En binding site (Figure 1A,B, construct *[15]*) resulted in a proportional increase of activity in the posterior wing compartment (75%, F(1,124) = 77.8, p=8.8818e-15). The log([*15]*/*[+]*) image (Figure 4) shows that mutant *[15]* proportionally affects the activity mostly in the posterior wing. The effect correlates with En distribution (Gompel et al., 2005) and is consistent with the repressive effect of its TF. Interestingly, mutant *[16]* shows a very similar *logRatio* to that of *[15]*, albeit with only 25% increase in activity. Mutations that disrupted characterized Dll binding sites (Figure 1A,B, constructs *[0], [1], [7]* and *[9]*) resulted in strong reduction in reporter expression (90% F(1,74) = 143.3, p=0; 75%, F(1,78) = 109.3, p=2.2204e-16; 47%, F(1,107) = 75.4, p=4.8073e-14 and 39%, F(1,74) = 23.2, p=7.6363e-06, respectively; Figure 1–figure supplement 3), which we quantified here for the first time. The *logRatio* images for mutants *[0], [1]*, and to a lesser extent *[7]*, show a patterned decrease of activity in line with Dll distribution in the wing (Figure 4) (Arnoult et al., 2013), with a proportionally stronger loss of activity toward the distal wing margin. This corroborates previous evidence that Dll binds to these sites. The respective *logRatio* images for segments *[0]* and *[1]* correlate with levels of Dll across the wing. This suggests that these sites individually integrate mostly Dll information, and do so in a near-linear fashion. Site *[9]* producing a relatively different picture with areas showing over-expression is discussed below. Mutations of Dll sites, however, clearly have non-additive effects, as mutants *[0], [1], [7]* and *[9]* result in a decrease of activity levels by 90%, 75%, 47% and 39% compared to *[+]*, respectively. This non-additivity could be explained by strong cooperative binding of Dll at these sites, or alternatively by considering that these Dll TFBS are interacting with other activating sites in the sequence.

**Figure 4.**
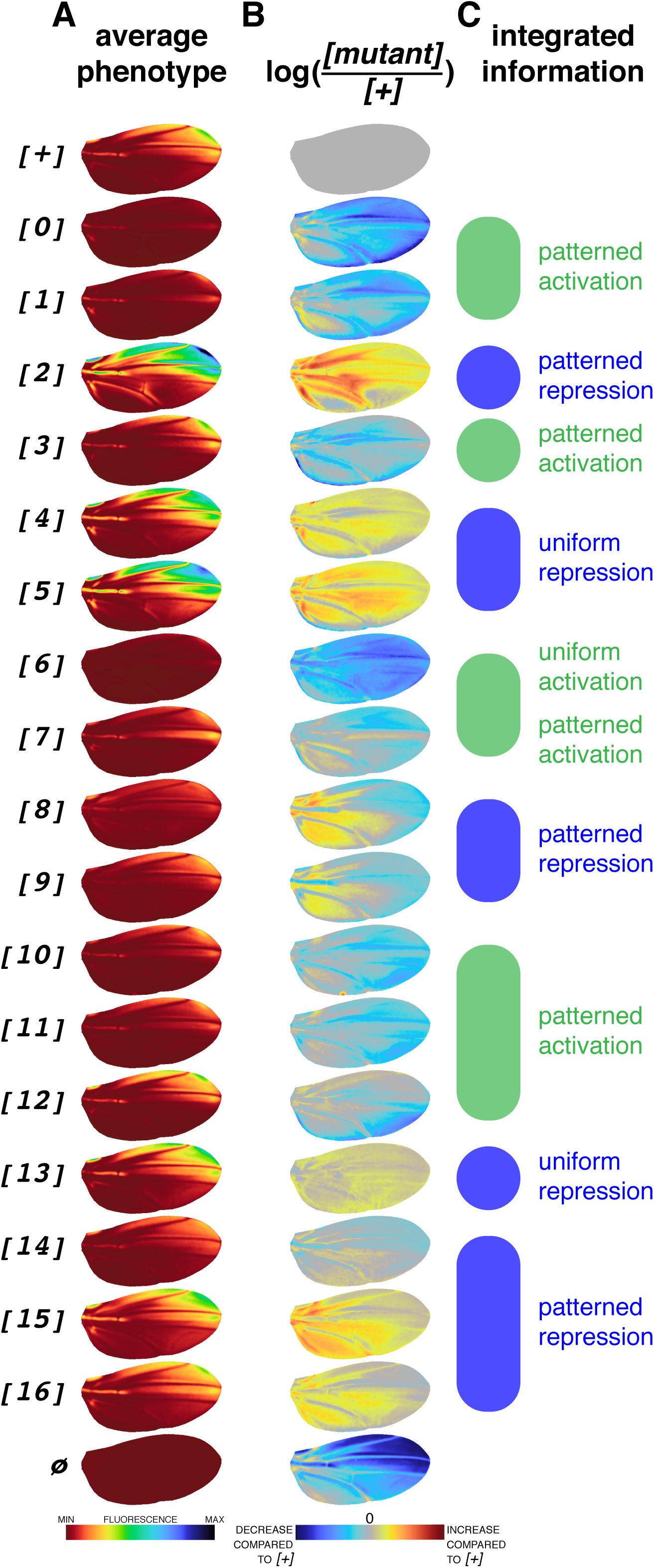
*trans*-regulatory integration along the *spot*^*196*^ sequence. (**A**) Average phenotypes reproduced from Figure 1B. (**B**) *logRatio* images (log(*[mutant]*/*[+]* for intensity values of each pixel of registered wing images) reveal what spatial information is integrated by each position along the enhancer sequence. For instance, a blue region on an image indicates that the enhancer position contains information for activation in this region. When mutated, this enhancer position results in lower activity than *[+]* in this region of the wing. Note that *logRatio* illustrates local changes between *[+]* and mutants far better than image differences (Figure 4–figure supplement 1) in regions of relatively low activity. (**C**) Summary of spatial information integrated along the enhancer sequence.

In addition, we noted that despite mutating a Dll TFBS, mutant *[9]* showed a substantially different *logRatio* than *[0]* and *[1]* but similar to *[8]*, with a repressing activity in the posterior wing compartment, proximally, and a distal activation (Figure 4B). This dual effect could be explained by the disruption of the Dll site along with a distinct TFBS for a posterior repressor. Alternatively, a single TFBS could be used by different TFs with opposite activities. In this regard, we note that the homeodomain of Dll and En have similar binding motifs (Zhu et al., 2011) and could both bind the Dll TFBS disrupted by *[9]* (and possibly *[8]*). The posterior repression of En and distal activation of Dll seems compatible with this hypothesis.

### Unraveling unexpected *trans*-regulatory integration along the *spot*^*196*^ sequence

We next analyzed the information integrated in other segments. In our previous dissections, not only did we overlook these segments, but we also did not expect them to affect the enhancer activity. Indeed, the combination of activation by Dll and posterior repression by En seemed sufficient to explain the spatial activity of *spot*^*196*^ without invoking additional regulators. Apart from the known Dll and En TFBSs, the enhancer scan of Figure 2 identified several segments with strong quantitative effects on the regulatory activity. Between the two pairs of Dll TFBSs, we found an alternation of activating sites (*[3]*and *[6]*, reducing overall levels by 36% (F(1,69) = 17.6, p=7.8336e-05) and 93% (F(1,98) = 284.9, p=0) compared to *[+]*, respectively), and strong repressing sites (*[2], [4]* and *[5]*, with an overall level increase of 3.2 folds (F(1,72) = 511.5, p=0), 1.9 folds (F(1,85) = 103.2, p=2.2204e-16) and 2.7 folds (F(1,82) = 426.5, p=0) compared to *[+]*, respectively). Construct *[3]* proportionally decreases the expression mostly around wing veins (Figure 4B), suggesting that this segment integrates information from an activator of the vein regions. We had found a similar activity for this region of *yellow* from another species, *D. pseudoobscura*, where no other wing activity concealed it (Gompel et al., 2005). Interestingly, the *logRatio* of mutant *[6]*, with a stronger, more uniform effect than for the other mutants that repress the activity, suggests a different *trans*-regulatory integration than Dll sites. In a separate study (Xin et al., 2020), we have further explored the functional role of segment *[6]* and have shown that this site regulates the chromatin state of the enhancer. Regarding segments with a repressive effect, mutants *[4]* and *[5]* result in a fairly uniform relative increase in expression, different from the activity of *[2]*, indicating that the information integrated by these two regions (*[2] vs. [4]* and *[5]*) likely involves different TFs. Together, three segments, *[6], [0]* and *[1]* (the last two containing previously known Dll binding sites), decrease the activity levels by 75% or more. Finding additional strong repressive sites (*[2], [4], [5]*) with a global effect on the enhancer activity across the wing is also unexpected.

The analysis revealed another activating stretch of sequence, between 116-137 bp, as mutating segments *[10]* and *[11]* decreased activity by 56% relative to *[+]* and showed very similar *logRatios*. Mutant *[12]* showed a mixed effect, with practically, in absolute terms, no effect in anterior distal wing quadrant. Finally, segments *[13], [14]*, and *[15]* showed a succession of repressing and activating sites, as we have seen for segments *[2]* - *[6]*, although with a lower amplitude. Mutant *[13]* caused an overall increase in activity (1.4 fold relative to *[+]*) with, proportionally, a uniform effect across the wing (*logRatio*). By contrast, mutant *[14]* decreased the overall activity by 36% with a *logRatio* indicating an activating effect in the spot region, and a repressive effect in the proximal part of the posterior wing compartment, similarly to mutants *[8]* and *[9]* but with lesser effects.

Together this first dissection, focusing on the necessity of segments for the enhancer activity at the scale of a TFBS, suggested a very different regulatory logic for the *spot*^*196*^ than previously described (Arnoult et al., 2013; Gompel et al., 2005). The non-additivity of effects at Dll binding sites, three repressing and four activating novel segments distributed in alternation along the enhancer, and the variety of their effects pointed to a more complex logic, involving more (possibly 6 to 8) factors than just Dll and En. We resorted to a different approach to further probe the regulatory logic of *spot*^*196*^.

### An interplay of activating and repressing inputs produces a spatial pattern of enhancer activity

The first series of mutations informed us on the contribution of the different elementary components of the *spot*^*196*^ enhancer sequence to its regulatory activity. Yet, it failed to explain how these components integrated by each segment interact to produce the enhancer activity. To unravel the regulatory logic of this enhancer, it is required to understand which segments are sufficient to drive expression, but also how elementary components underlying the regulatory logic influence each other. To evaluate the sufficiency of, and interactions between, different segments, would require to test all possible combinations of mutated segments, namely a combinatorial dissection. Doing this at the same segment resolution as above is unrealistic, as the number of constructs grows with each permutation. Instead, we used three sequence blocks of comparable sizes in the *spot*^*196*^ enhancer, *A, B* and *C*, defined arbitrarily (Figure 5A), and produced constructs where selected blocks were replaced by randomized sequence (noted “*-*”). This second series, therefore, consists of nine constructs, including all combinations of one, two or three randomized blocks, a wild type *[ABC]* (which has strictly the same sequence as *[+]* from the first series) and a fully randomized sequence, *[---]*.

**Figure 5.**
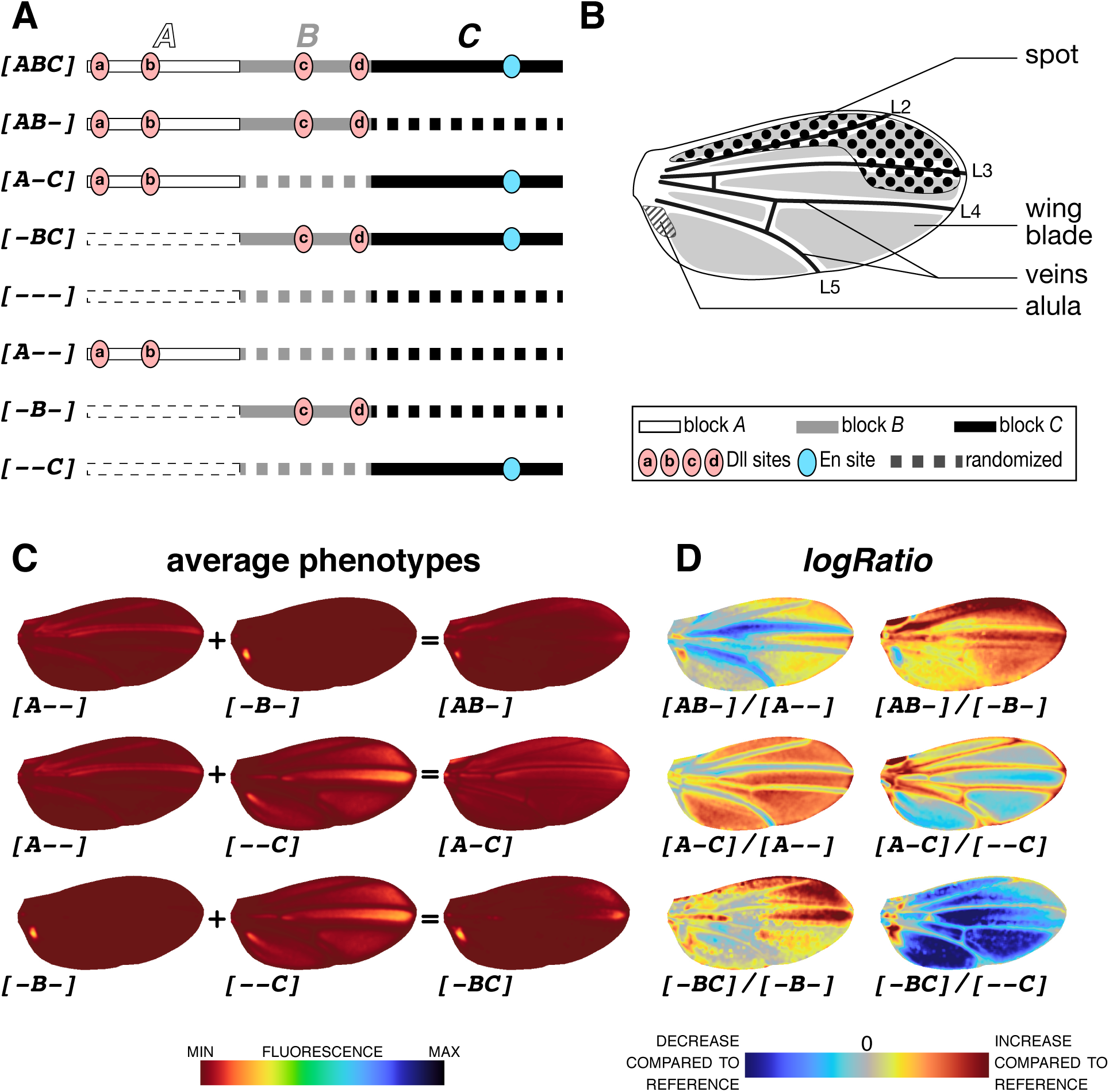
Regulatory interactions in the *spot*^*196*^ sequence. (**A**) Schematics of constructs with block randomizations. *spot*^*196*^ sequence was arbitrarily divided into 3 blocks (A: 63 bp; B: 54 bp; C: 79 bp). In each construct, the sequence of one, two or all 3 blocks was randomized. (**B**) Terminology for parts of the wing where constructs from (**A**) drive reporter expression. (**C**) Average phenotypes resulting from constructs in (**A**). Constructs where single blocks remain indicate the sufficiency of these blocks to promote wing activity: *A* in the veins, *B* in the alula and *C* at high levels across the wing blade. Constructs with two non-randomized blocks show the effect of one block on the other. For instance, *B* is sufficient to suppress the wing blade activation promoted by *C*, as seen by comparing *[-B-], [--C]* and *[-BC]*. Colormap of average phenotypes normalized for all constructs of the block series, including block permutations of Figure 6B. (**D**) Block interactions is best visualized with *logRatio* images of constructs phenotypes shown in (**C**). For each *logRatio*, the denominator is the reference construct and the image shows on a logarithmic scale how much the construct in the numerator changes compared to this reference. For instance, log(*[-BC]*/*[--C]*) shows the effect of *B* on *C*, a global repression, except in the spot region. Colormap indicates an increase or a decrease of activity compared to the reference (denominator). For an overview of all comparisons, particularly the relative contribution of each block to the entire enhancer activity, see Figure 5–figure supplement 2C-F.

With these constructs, we can track which segments, identified in the first series as necessary for activation in the context of the whole *spot*^*196*^, are also sufficient to drive activity (Figure 5–figure supplement 1; see Figure 2 for the correspondence between the two series of mutations). Of the three blocks (constructs *[A--], [-B-]* and *[--C]*), only the block *C* is sufficient to produce activity levels comparable to those of the wild-type *spot*^*196*^ in the wing blade, although with a different pattern from *[ABC]* (Figure 5–figure supplement 2A-C). Reciprocally, randomizing block *C* (construct *[AB-]*) results in a uniform collapse of the activity (Figure 5–figure supplement 2A-C). We concluded that the sequence of block *C* contains information necessary and sufficient to drive high levels of activity. This is particularly interesting because *C* does not contain previously identified Dll TFBSs, or strong activating segments. By contrast, blocks *A* and *B*, although they each contain two Dll sites, do not drive wing blade expression. The activating segments in block *C* revealed in the first dissection, particularly segments *[10]* and *[11]* are therefore candidates to drive the main activity of the *spot*^*196*^, in the context of these reporter constructs.

Block *A* alone (*[A--]*) produces high levels of expression in the veins (Figure 5–figure supplement 2A-C). Combined with block *C* (construct *[A-C]*), it also increases the vein expression compared to *C* alone. We concluded that *A* is sufficient to drive expression in the veins. Segment *[3]*, which proportionally decreased the activity mostly in the veins could therefore be the necessary counterpart for this activation.

Block *B* alone drives expression only near the wing hinge, in a region called the alula (*[-B-]*, Figure 5B-D). The first dissection series, however, did not identify a mutated segment within block *B* that affected specifically the alula.

The necessity of Dll binding sites (in segments *[0], [1], [7]* and *[9]*) and of segment *[6]*, and their insufficiency to drive activity in the wing blade in the context of block *A* alone, block *B* alone, or blocks *A* and *B* combined, indicate that these strong activating sites function in fact as permissive sites. We next focused on understanding the interplay between repressing and activating sites, to shed light on how the *spot*^*196*^ patterning information is built. In the first series of constructs, we identified several strong repressing segments in block *A* (*[2]* and *[4]*) and block *B* (*[5]*). Using sufficiency reasoning with the second series of constructs, we further investigated how these inputs interacted with other parts of the enhancer (Figure 5). Such interactions are best visualized with *logRatios*, comparing this time double-block constructs to single-block constructs used as references (Figure 5D and Figure 5–figure supplement 2D-F). Block *B* has a strong repressive effect on block *C* throughout the wing, except at the anterior distal tip, where *C* activity is nearly unchanged (log(*[-BC]*/*[--C]*), Figure 5D). Likewise, log(*[AB-]*/*[A--]*) shows that *B* also represses the vein expression driven by *A*. Similarly, block *A* represses the *C* activity across the wing blade, except in the spot region log(*[A-C]*/*[--C]*). We have seen above that blocks *A* and *B* both contain strong repressing segments, but also known Dll TFBSs. Because both *A* and *B* show a repressive effect on block *C*, except in the spot region, we submit that the apparent patterned activation by Dll may in fact result from its repressive effect on direct repressors of activity, mostly at the wing tip. This indirect activation model would explain the non-additivity of the individual Dll binding sites observed in the first construct series and why grafting Dll TFBSs on a naïve DNA sequence is not sufficient to create a wing spot pattern.

Together, these results outline an unexpectedly complex regulatory logic contrasting with the simple model we had initially proposed (Arnoult et al., 2013; Gompel et al., 2005). In a model of regulatory logic summarized in Figure 7, we find that at least two direct activators, a vein activator (*[3]*) and a wing blade activator (potentially *[10]* and *[11]*) promote activity throughout the wing, which is in turn repressed everywhere (*[2], [4]* and *[5]*) except where Dll is present. In addition, En represses the action of Dll in the posterior wing compartment. The exact role of the permissive segment *[6]* is still unclear, as it showed a strong effect, but is not necessary for the activity driven by block *C*. It may be required to grant some TFs access to the enhancer (Xin et al., 2020).

**Figure 6.**
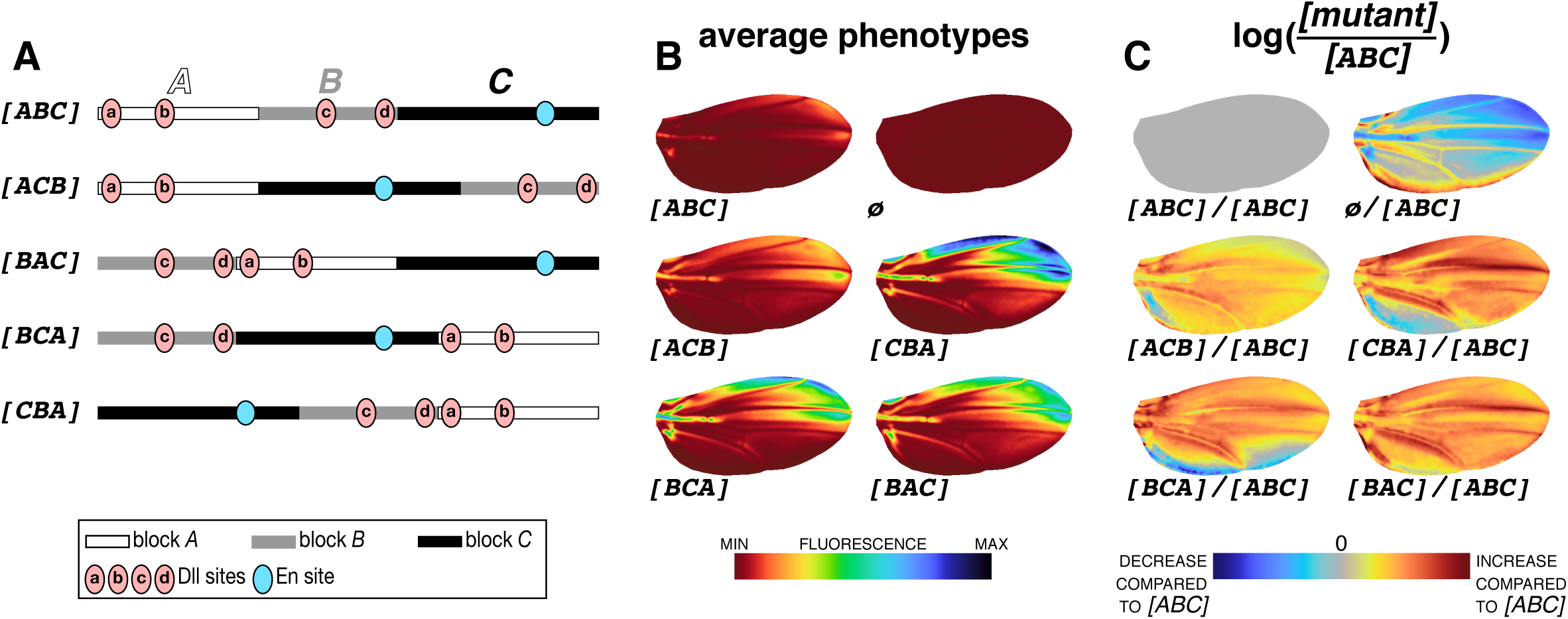
Block permutations scale the activity of the *spot*^*196*^ enhancer. (**A**) Schematics of constructs with block permutations. In this series, the same blocks of sequences as in Figure 5A were permutated. (**B**) Average phenotypes resulting from constructs in (**A**). Colormap of average phenotypes normalized for all constructs of the block series, including block randomizations of Figure 5C and Figure 5–figure supplement 2B. (**C**) Average phenotypes in (**B**) compared to the average phenotype of the wild type *[ABC]* (*logRatio*). Note that, in contrast to constructs with randomized blocks (Figure 5), constructs with block permutations results in near-uniform changes of activity across the wing. Colormap indicates an increase or a decrease of activity compared to the wild-type enhancer *[ABC]*.

**Figure 7.**
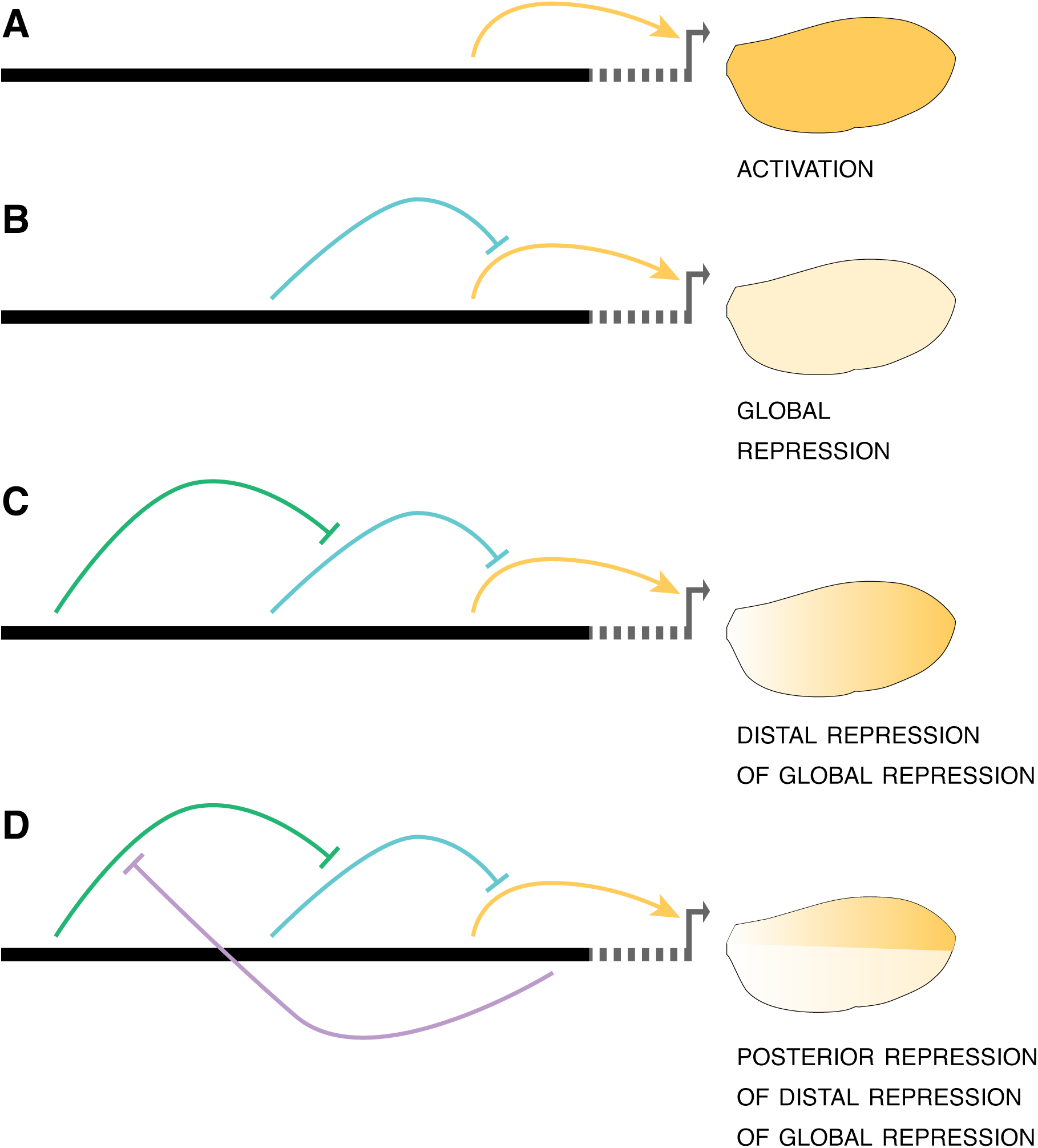
A model of the regulatory logic governing the *spot*^*196*^ enhancer. (**A**)-(**D**) The schematics shows step by step how regulatory information and interactions integrated along the enhancer sequence produce a spatial pattern of activity. (**A**) three independent inputs, respectively in blocks *A, B* and *C*, promote activity (arrows) in the wing veins, the alula and the wing blade, as illustrated with average phenotypes of constructs *[A--], [-B-]* and *[--C]*, respectively. Note that activity levels in the wing blade, stemming from block *C*, match the final levels of the *spot*^*196*^ enhancer activity in the spot region. (**B**) a first set of repressive inputs suppress activity in the wing blade (stemming from blocks *A* and *B*) and the veins (stemming from blocks *B*). The overall combined output of the initial activation and the global repressive inputs is a near complete loss of activity, except in the alula. (**C**) A second set of repressive inputs, whose action is localized in the distal wing region, counters the global repression, thereby carving a pattern of distal activity promoted by block *C*. (**D**) The distal activity is repressed in the posterior wing compartment, likely through the repressive action of Engrailed, resulting in a final pattern of activity in the spot region.

### Sequence reorganization affects activity levels of the *spot*^*196*^ enhancer, not its spatial output

In a final series of experiments, we wondered whether the complex regulatory architecture uncovered by the first two mutant series was sensitive to the organization of the inputs. To test the effect of changes in the organization of enhancer logical elements, we introduced new constructs with permutations of blocks *A, B* and *C* (Figure 6A). These permutations preserve the entire regulatory content of the enhancer, except at the junction of adjacent blocks where regulatory information may be lost or created. All permutations that we have tested (4 out of 5 possible permutations) drive significantly higher levels of expression than the wild type *[ABC]* (*[ACB]*: 2.9 folds (F(1,98) = 191.8, p=0); *[BAC]*: 6 folds (F(1,93) = 589.1, p=0); *[BCA]*: 5.8 fold (F(1,93) = 589.1, p=0); *[CBA]*: 8.4 folds (F(1,93) = 1664.2, p=0); Figure 6B), yet with minor effects on the activity distribution proportionally to the wild type (Figure 6C). We concluded from these experiments that, in terms of pattern, the regulatory output is generally resilient to large-scale rearrangements. As long as all inputs are present in the sequence, the spatial activity is deployed in a similar pattern, yet its quantitative activity is strongly modulated. Because they have little influence on the activity pattern, the rearrangements may not change the nature of the interactions within the enhancer or with the core promoter. Although we would need to challenge this conclusion with additional constructs and blocks with different breakpoints, we speculate that, molecularly, the block randomization perturbates the action of some of the uniformly repressing elements. It highlights the robustness of the enhancer logic to produce a given patterned activity.

## Discussion

With this work, we have set to decipher the regulatory logic of an enhancer, *spot*^*196*^. The view point presented here, devolving from systematic necessity and sufficiency analysis in *cis*, is the information that the enhancer integrates along its sequence. Combined with the quantitative measurement of enhancer activity in a tissue, the wing, this information reveals the enhancer regulatory logic and how it reads the wing *trans*-regulatory environment to encode a spatial pattern.

### Quantitative assessment of gene expression to decipher regulatory logic

Before discussing this regulatory logic, it is relevant to underscore the importance of considering gene expression comprehensively and quantitatively in analyzing developmental processes. The spectacular progress of developmental biology in the last four decades owes much to the careful analysis of phenotypes. Allelic mutant series with varying levels of gene expression, resulting in gradually stronger phenotypes, imposed the notion that development is a quantitative process. Geneticists and cell biologists have formalized developmental processes in terms of spatial and temporal patterns of gene expression (Carroll, 1998), whereas the quantitative dimension is often left out. The overall construction and architecture of an embryo undoubtedly depends on the time and location of gene expression, but the amount of each gene product tunes the properties of many traits, from the position of cell population boundaries to the physical properties of an organ. Moreover, limiting the assessment of gene expression to its qualitative aspects (a spatial pattern assessed visually, not quantitatively) biases the analyses to the modifications that produce major effects, potentially hindering the discovery of essential or important determinants, as shown in the present study. The limited number of developmental biology studies using thorough quantitative measures of gene expression in tissues is for an obvious reason. It is much more difficult and requires new technical and conceptual advances to describe morphological phenotypes comprehensively and quantitatively. The advent of genome-wide methods –omics– to reconstruct spatio-temporal gene expression represents of course a major step in that respect, but, even with single-cell omics, assigning the precise distribution of gene transcript to cells is difficult and indirect (Nitzan et al., 2019). Several studies have successfully measured direct gene expression quantitatively during development and linked it to morphology. These attempts range from modeling a monotone gradient (Gregor et al., 2007), to quantifying variation in gene expression patterns (Crocker and Stern, 2017; Fowlkes et al., 2008; Knowles, 2012; Martinez-Abadias et al., 2018). To our knowledge, however, very few (Bentovim et al., 2017; Park et al., 2019) used comprehensive descriptors to thoroughly extract more information than possible from qualitative approaches, or exploited the statistical power offered by such descriptors. This prompted us to implement a generic quantitative framework to measure and compare activity levels of the *spot*^*196*^ enhancer across the wing among genotypes, and use powerful statistical tools to test the significance of the differences.

### Regulatory necessity and regulatory sufficiency

The strength of our arguments also stems from the introduction of two complementary aspects of the method: one to combine the assessment of necessity and sufficiency of regulatory information in our analysis (discussed in this section) and another to compare the activity of enhancer variants (*logRatio*, discussed in the next section).

When dissecting a regulatory element, it is straightforward to assess the necessity of a TFBS or any stretch of sequence to the activity, by introducing mutations. It is generally more difficult to assess whether the same sequence is sufficient to promote regulatory activity at all and most enhancer dissections are focusing on necessity analysis (see for instance(Arnosti et al., 1996; Arnoult et al., 2013; Bertrand et al., 2003; Farley et al., 2016; Gompel et al., 2005; Park et al., 2019; Swanson et al., 2010; Swanson et al., 2011; Thanos and Maniatis, 1995)). Yet, our study clearly shows that to decipher regulatory logic, and eventually design synthetic enhancers, understanding which regulatory components are sufficient to build an enhancer activity is key.

### A visual tool to compare spatial activities driven by enhancer variants

We introduced a new representation to compare activities between enhancer variants, typically a wild type and a mutant. Proportional effects, or local fold changes, as revealed by *logRatio* produce representations that are independent from the distribution of the reference activity. They also reflect better the distribution of factors in *trans* and their variations as seen by the enhancer (here, across the wing) than differential comparisons. Indeed, differential comparisons are dominated by regions of high activities and thereby focusing our attention to the regions of high variation of activity. By contrast, along with a large number of samples and precise quantification, *logRatios* reveal strong effects in regions of low activity that would hardly be visible using differential comparisons, highlighting some cryptic components of the regulatory logic. When additional knowledge about TFBSs and TF distribution will become available, they will also inform us on the contribution of the TF in the regulatory logic. In this respect, the introduction of *logRatios* in our analysis has proven useful and could be adapted to any system where image alignment is possible, such as *Drosophila* blastoderm embryos (Fowlkes et al., 2008), or developing mouse limbs (Martinez-Abadias et al., 2016).

### A-tracts did not disrupt major effect of TF-TF interactions

A-tracts are known to change local conformational properties of DNA. As such, our A-tract mutations could influence the regulatory logic not only by directly disrupting the information contained in the sequence they replaced, but also indirectly, by introducing more changes than wanted. As an alternative, sequence randomization, however, is more likely to create spurious TFBSs, which is difficult to control for, especially if all the determinants of the enhancer activity are not known. The possible occurrence of undesired and undetected TFBSs would have biased our interpretation of the effect of individual segments, and consequently, of the regulatory logic of the enhancer. The chance that A-tracts introduce new TFBSs in the enhancer sequence is quite low compared to sequence randomization, which is why we favored this mutational approach. Yet, A-tracts can modify various physical properties of the DNA molecule, and in turn, influence interactions between TFs binding the enhancer. The disruption of a TF-TF interaction due to the introduction of an A-tract between two TFBSs (Fig. 3B) would be revealed if mutating a particular segment would have an effect similar to the effect of mutating immediately adjacent flanking segments. We note, however, that we do not have such situation in our dataset. This suggests that the A-tracts we introduced, if anything, only mildly altered TF-TF interactions through changes in the physical properties of *spot*^*196*^. Instead, we think that the effects of A-tract mutations are mostly due to disrupted TFBSs along the enhancer sequence.

### The regulatory logic underlying *spot*^*196*^ enhancer activity

The main finding of our study is that the *spot*^*196*^ enhancer integrates 6 to 8 distinct regulatory inputs, with multiple layers of cross-interactions (Figure 7). We had previously proposed that the spot pattern resulted from the integration of only two spatial regulators, the activator Dll, and the repressor En (Arnoult et al., 2013; Gompel et al., 2005). A logical analysis of systematic mutations along the enhancer, gives a different status to these factors. The main levels of *spot*^*196*^ activity across the wing blade seem to result mostly from two unknown activators, one promoting a relatively uniform expression in the wing blade, and another along the veins (Figure 7A). This activation is in turn globally repressed throughout the wing by an unknown repressor whose action masks that of the global activator (Figure 7B). Upon this first two regulatory layers, the actual spot pattern of activity is carved by two local repression. A distal repression counteracts the effect of the global repressor in the distal region of the wing (Figure 7C) but the spatial range of this repression is limited to the anterior wing compartment by another repressor acting across the posterior wing compartment (Figure 7D). The former local repression could be mediated by Dll itself, a hypothesis compatible with the non-additive effects of Dll TFBS mutations, while the latter is almost certainly due to En. Thus, the pattern of activity results not so much from local activation but from a complex, multilayered interplay of repressors.

One would expect this complex set of interactions between TFs that bind along the enhancer sequence to be vulnerable to sequence reorganization. We find surprising that shuffling blocks of sequence resulted in dramatic changes in activity levels with little effect on the activity pattern. Similarly, many of the mutations still produced a pattern of activity quite similar to the one of *[+]*. This suggests that the exact organization of the different inputs, and the absence of some of these inputs, do not affect the TF-enhancer and TF-TF interactions required for a patterned activity, which here translates mainly to the role of Dll in repressing global repressors, and the repressing role of En. The frequency of these interactions, or the interactions with the core promoter, may, however, change significantly upon sequence modifications, impacting transcription rate. In other words, the regulatory logic described above is robust to changes for the production of a spatial pattern, but less so for the tuning of enhancer activity levels.

The evolutionary steps of the emergence of *spot*^*196*^ perhaps reflect in the regulatory logic of this enhancer. The *spot*^*196*^ element evolved from the co-option of a pre-existing *wing blade* enhancer (Gompel et al., 2005). The sequences of this ancestral *wing blade* enhancer and the evolutionary-derived *spot*^*196*^ overlap and share at least one common input (Xin et al., 2020). This perspective is consistent with the idea that a novel pattern emerged by the progressive evolution of multiple repressive layers carving a spot pattern from a uniform regulatory activity in the wing blade. To further deconstruct the regulatory logic governing the *spot*^*196*^ enhancer and its evolution, one first task will be to investigate how some of the mutations we introduced impact the activity of a broader fragment containing the entire *spot* activity (and the *wing blade* enhancer), closer to the native context of this enhancer. Another challenging step will be to identify the direct inputs integrated along its sequence. It will also be necessary to characterize their biochemical interactions with DNA and with one another. Ultimately, to fully grasp the enhancer logic will mean to be able to recreate these interactions in a functional synthetic regulatory element.

## Materials and Methods

### Fly husbandry

Our *Drosophila melanogaster* stocks were maintained on standard cornmeal medium at 25°C with a 12:12 day-night light cycle.

### Transgenesis

All reporter constructs were injected as in (Arnoult et al., 2013). We used ϕC31-mediated transgenesis (Groth et al., 2004) and integrated all constructs at the genomic *attP* site *VK00016* (Venken et al., 2006) on chromosome 2. All transgenic lines were genotyped to ascertain that the enhancer sequence was correct.

### Molecular biology

All 196 bp constructs derived from the *D. biarmipes spot*^*196*^ sequence were synthetized *in vitro* by a Biotech company (Integrated DNA Technologies, Coralville, United States, Cat. #121416). Figure 1–figure supplement 1 provides a list of all constructs and their sequences. Each construct was cloned by In-Fusion (Takara, Mountain View, United States) in our pRedSA vector (a custom version of the transformation vector pRed H-Stinger (Barolo et al., 2004) with a 284 bp *attB* site for ϕC31-mediated transgenesis (Groth et al., 2004) cloned at the AvrII site of pRed H-Stinger). All constructs in Figure 1 were cloned by cutting pRedSA with Kpn I and Nhe I, and using the following homology arms for In-Fusion cloning: 5’-GAGCCCGGGCGAATT-3’ and 5’-GATCCCTCGAGGAGC**-**3’. Likewise, constructs in Figure 5 were cloned by cutting pRedSA with BamH I and EcoR I, and using the following homology arms for In-Fusion cloning: 5’-GAGCCCGGGCGAATT-3’ and 5’-GATCCCTCGAGGAGC**-**3’.

### Wing preparation and imaging

All transgenic wings imaged in this study were homozygous for the reporter construct. Males were selected at emergence from pupa, a stage that we call “post-emergence”, when their wings are unfolded but still slightly curled. When flies were massively emerging from an amplified stock, we collected every 10 minutes and froze staged flies at −20°C until we had reached a sufficient number of flies. In any case, staged flies were processed after a maximum of 48 hours at - 20°C. We dissected a single wing per male. Upon dissection, wings were immediately mounted onto a microscope slide coated with transparent glue (see below), and fixed for 1 hour at room temperature in 4% paraformaldehyde diluted in phosphate buffer saline 1% Triton X-100 (PBST). Slides with mounted wings were then rinsed in PBST and kept in a PBST bath at 4°C until the next day. Slides were then removed from PBST and the wings covered with Vectashield (Vector Laboratories, Burlingame, United States). The samples were then covered with a coverslip. Preparations were stored for a maximum of 48 hours at 4°C until image acquisition.

The glue-coated slides were prepared immediately before wing mounting by dissolving adhesive tape (Tesa brand, tesafilm®, ref. 57912) in heptane (2 rolls in 100 ml heptane), and spreading a thin layer of this solution onto a clean microscope slide. Once the heptane had evaporated (under a fume hood), the slide was ready for wing mounting. All wing images were acquired as 16-bit images on Ti2-Eclipse Nikon microscope equipped with a Nikon 10x plan apochromatic lens (N.A. 0.45; Nikon Corporation, Tokyo, Japan) and a pco.edge 5.5 Mpx sCMOS camera (PCO, Kelheim, Germany) under illumination from a Lumencor SOLA SE II light source (Lumencor, Beaverton, OR, USA). Each wing was imaged by tiling and stitching of several z-stacks (z-step = 4 µm) with 50% overlap between tiles. Each image comprises a fluorescent (ET-DSRed filter cube, Chroma Technology Corporation, Bellows Falls, VT, USA) and a bright field channel (acquired using flat field correction from the Nikon NIS-Elements software throughout), the latter being used for later image alignment. To ensure that fluorescence measurements are comparable between imaging sessions, we have used identical settings for the fluorescence light source (100 % output), light path and camera (20 ms exposure time, no active shutter) to achieve comparable fluorescence excitation.

### z-Projection

Stitched 3D stacks were projected to 2D images for subsequent analysis. The local sharpness average of the bright-field channel was computed for each pixel position in each z-slice and an index of the slice with the maximum sharpness was recorded and smoothed with a Gaussian kernel (sigma = 5 px). Both bright-field and fluorescent 2D images were reconstituted by taking the value of the sharpest slice for each pixel.

### Image alignment

Wing images were aligned using the veins as a reference. 14 landmarks placed on vein intersections and end points, and 26 sliding landmarks equally spaced along the veins were placed on bright field images using a semi-automatized pipeline. Landmark coordinates on the image were then used to warp bright field and fluorescent images to match the landmarks of an arbitrarily chosen reference wing by the thin plate spline interpolation (Hutchinson, 1995). All wings were then in the same coordinate system, defined by their venation.

### Fluorescent signal description

A transgenic line with an empty reporter vector (*ø*) was used as a proxy to measure noise and tissue autofluorescence. The median raw fluorescent image was computed across all *ø* images and used to remove autofluorescence, subtracted from all raw images before the following steps. All variation of fluorescence below the median *ø* value was discarded. The DsRed reporter signal was mostly localized in the cell nuclei. We measured the local average fluorescent levels by smoothing fluorescence intensity, through a Gaussian filter (sigma = 8 px) on the raw 2D fluorescent signal. The sigma corresponded roughly to 2 times the distance between adjacent nuclei. To lower the memory requirement, images were then subsampled by a factor of 2. We used the 89735 pixels inside the wings as descriptors of the phenotype for all subsequence analyses.

### Average phenotypes, differences, logRatio colormaps and normalization

Average reporter expression phenotypes were computed as the average smoothed fluorescence intensity at every pixel among all individuals in a given group (tens of individuals from the same transgenic line). The difference between groups was computed as the pixel-wise difference between the average of the groups (Figure 4–figure supplement 1). *logRatio* between two constructs represents the fold change of a phenotype relatively to another and is calculated as the pixel-wise logarithm of the ratio between the two phenotypes. Averages, difference, and *logRatio* images were represented using colors equally spaced in CIELAB perceptual color space ((CIE), 2018). With these colormaps the perceived difference in colors corresponds to the actual difference in signal. Colormaps were spread between the minimal and maximal signals across all averages for average phenotypes. Difference and *logRatio* spread between minus and plus represent the absolute value of all difference for the phenotype differences, grey colors meaning that the two compared phenotypes are equal.

### Mutation effect direction and intensity

We proposed to represent the necessity of a stretch of sequence along the enhancer with the activity levels of mutants of this stretch relatively to wild-type (*[+]*) activity. To summarize the overall effect of mutants (overexpression or underexpression), we measured the average level of activity across the whole wing relatively to that of *[+]*. The effect of a mutation is not strictly limited to the mutated bases, as they can also modify properties of DNA of flanking positions (Zhou et al., 2013). To take this effect into account and produce a more realistic and conservative estimation of necessity measure at each position, we weighted the phenotypic contribution of each mutant line to the measure by the strength of the changes they introduce to the DNA shape descriptors at this position. At each position, the phenotype of constructs not affecting the DNA shape descriptors compared to *[+]* were not considered. When two mutants modify the DNA shape descriptors at one position, typically near the junction of two adjacent mutations, the effect at this position was computed as the weighted average of the effect of the two mutants, where the weight is the extent of the DNA shape modification relatively to *[+]* sequence. DNA shape descriptors were computed by the R package DNAshapeR (Chiu et al., 2016).

### Principal component analysis (PCA), and difference significance

The intensity measure is an average of the overall and variable expression across the wing. Hence, mutations causing a different effect on the phenotype can have the same intensity value. To test whether mutant significantly differ from *[+]*, we used comprehensive and unbiased phenotype descriptors provided by principal component analysis (PCA), which removes correlation between pixel intensities and describe the variation in reporter gene expression. PCA was calculated on the matrix regrouping intensities of all pixels for every individual, of dimensions (n_individuals x n_pixels on the wing). The significance of the difference between two constructs considers the multivariate variation of the phenotypes, and is tested using MANOVA on all 5 first components explaining more than 0.5% of the total variance (Figure 2–figure supplement 1).

### Overall expression intensity and significance

The overall expression level was measured for each individual as the average intensity across the wing. This was used to test the significance of overall increase and decrease in expression levels relatively to the wild-type levels.

### DNA rigidity scores

A-tracts are runs of consecutive A/T bp without a TpA step. Stacking interactions and inter-bp hydrogen bonds in ApA (TpT) or ApT steps of A-tracts lead to conformational rigidity (Nelson et al., 1987). The length of an A-tract directly correlates with increased rigidity (Rohs et al., 2009). To parametrize DNA rigidity at nucleotide resolution, we used A-tract length as a metric. For each position in a given DNA sequence, we find the longest consecutive run of the form A_*n*_T_*m*_ that contains this position (with the requirement of *n≥*0, *m≥*0, and *n*+*m≥*2), and score DNA rigidity at that position using the length of this sub-sequence. For example, the sequence AATCGCAT will map to the scores 3,3,3,0,0,0,2,2 because AAT and AT are A-tracts of lengths 3 and 2 bp, respectively.

## Author contributions

Yann Le Poul: Conceptualization, Methodology, Software, Validation, Formal analysis, Data curation, Writing—original draft, Visualization

Yaqun Xin: Validation, Investigation, Formal analysis, Data curation

Liucong Ling: Investigation, Formal analysis

Bettina Mühling: Investigation

Rita Jaenichen: Investigation

David Hörl: Software, Data curation

David Bunk: Software, Data curation

Hartmann Harz: Methodology, Supervision

Heinrich Leonhardt: Supervision

Yingfei Wang: Methodology, Software, Formal analysis

Elena Osipova: Investigation

Mariam Museridze: Investigation, Formal analysis

Deepak Dharmadhikari: Investigation, Formal analysis

Eamonn Murphy: Investigation, Formal analysis

Remo Rohs: Methodology, Supervision, Funding acquisition

Stephan Preibisch: Software, Supervision, Funding acquisition

Benjamin Prud’homme: Conceptualization, Writing—original draft

Nicolas Gompel: Conceptualization, Validation, Writing—original draft, Visualization, Supervision, Project administration, Funding acquisition

## Acknowledgements

This work was supported by funds from the Ludwig Maximilian University of Munich, the Human Frontiers Science Program (Program Grant RGP0021/2018 to NG, SP and RR), the Deutsche Forschungsgemeinschaft (grants INST 86/1783-1 LAGG and GO 2495/5-1 to NG), the European Research Council under the European Union’s Seventh Framework Programme (FP/2007-2013 / ERC Grant Agreement n° 615789 to BP) and the National Institutes of Health (grant R35GM130376 to RR). YX was supported by a fellowship from the China Scholarship Council (fellowship 201506990003). LL was supported by a DFG fellowship through the Graduate School of Quantitative Biosciences Munich (QBM). MM and DD are recipients of fellowships from the German Academic Exchange Service (DAAD). EM was supported by the Amgen Scholar program of the LMU.

## Legend to figures

**Figure 1–figure supplement 1**. Constructs sequences.

**Figure 1–figure supplement 2.**
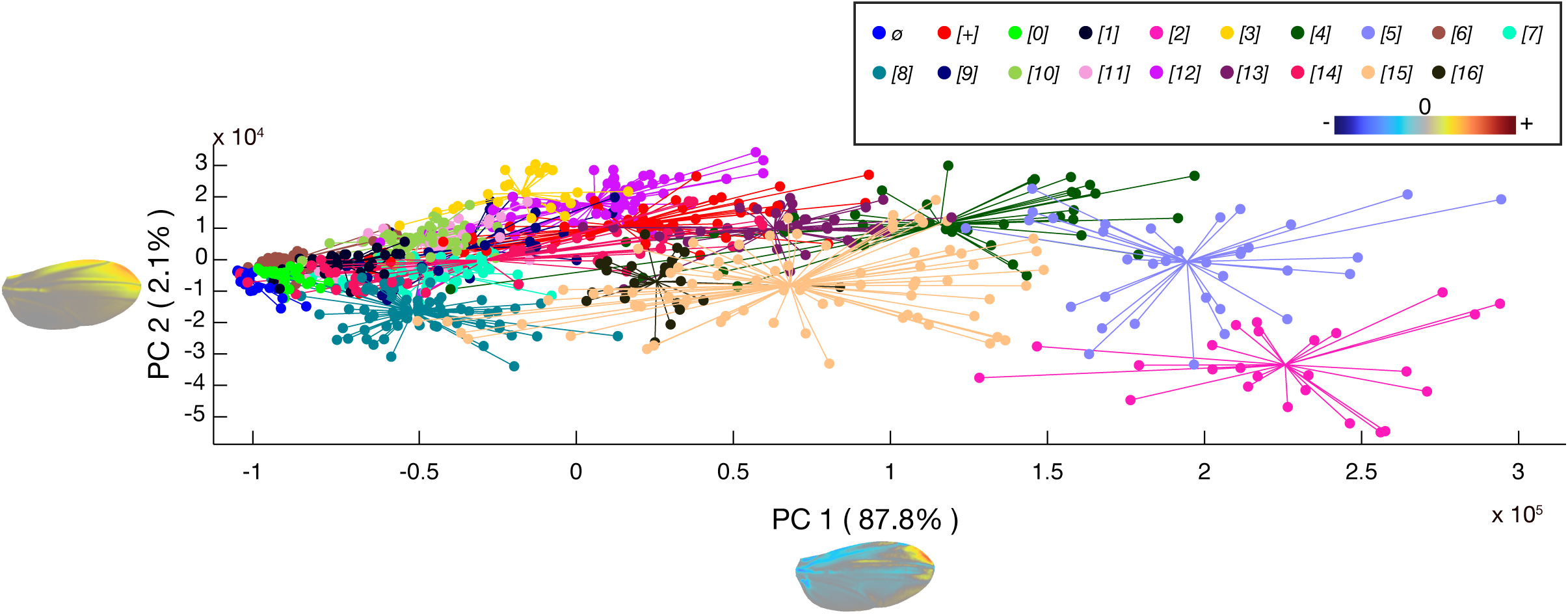
First two axes of variation in a principal component analysis of all individual wings used to generate the average reporter expression of Figure 1. Each wing is depicted by a colored dot, and each construct by a color. PC1 captures 87.8% of the variation and corresponds to overall changes in the activity of the *spot*^*196*^ CRE. PC2 captures 2.1% of the variation and appears to represent spatial difference in CRE activity between lines. The direction of variation along each principal component is represented on a wing with a colormap next to each axis.

**Figure 1–figure supplement 3**. Scores for the PCA shown in Figure 1–figure supplement 2.

**Figure 1–figure supplement 4**. Significance of difference in activity between pairs of groups, using the first 6 principal components.

**Figure 1–figure supplement 5**. Number of individuals analyzed for each construct in this study.

**Figure 2–figure supplement 1**. Significance of the difference in average expression levels among constructs of the first mutant series (*[0]*-*[16]*).

**Figure 4–figure supplement 1.**
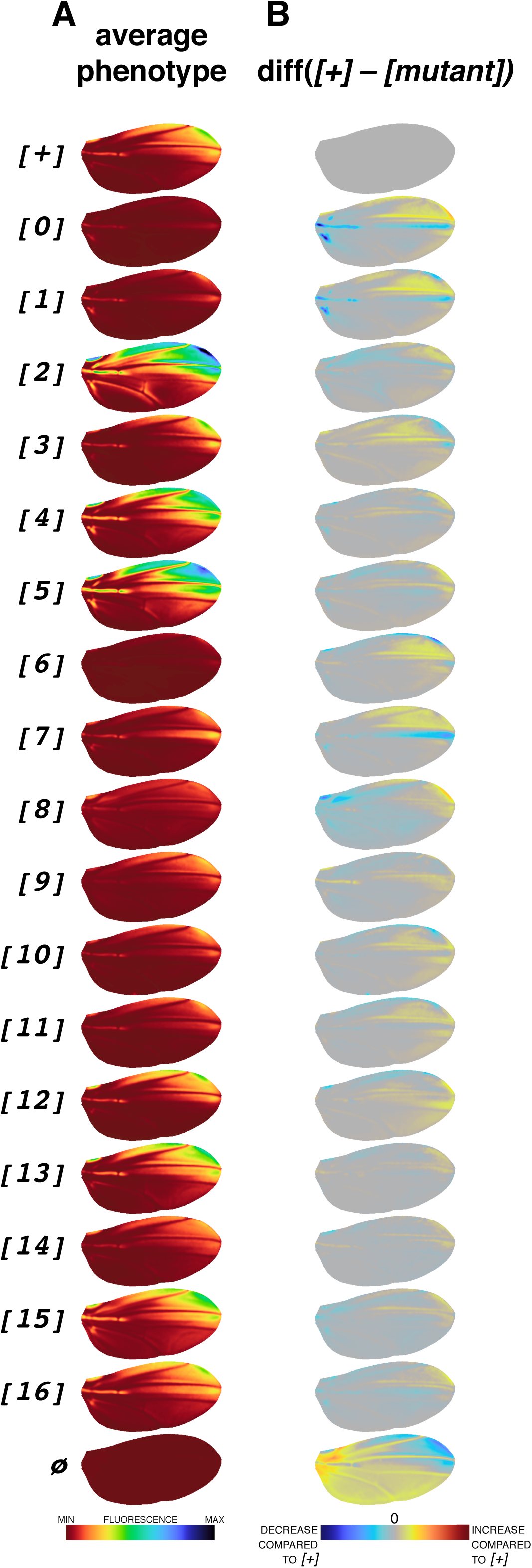
Pattern changes between wild-type and mutant *spot*^*196*^ constructs. (**A**) Average phenotypes reproduced from Figure 1B. (**B**) difference images (*[+]* – *[mutant]*) for intensity values of each pixel of registered wing images) highlight changes in the distribution of the enhancer activity across the wing. Note that this operation introduces a visual bias towards changes in region of high expression, contrasting with *logRatio* images of Figure 4.

**Figure 5–figure supplement 1.**
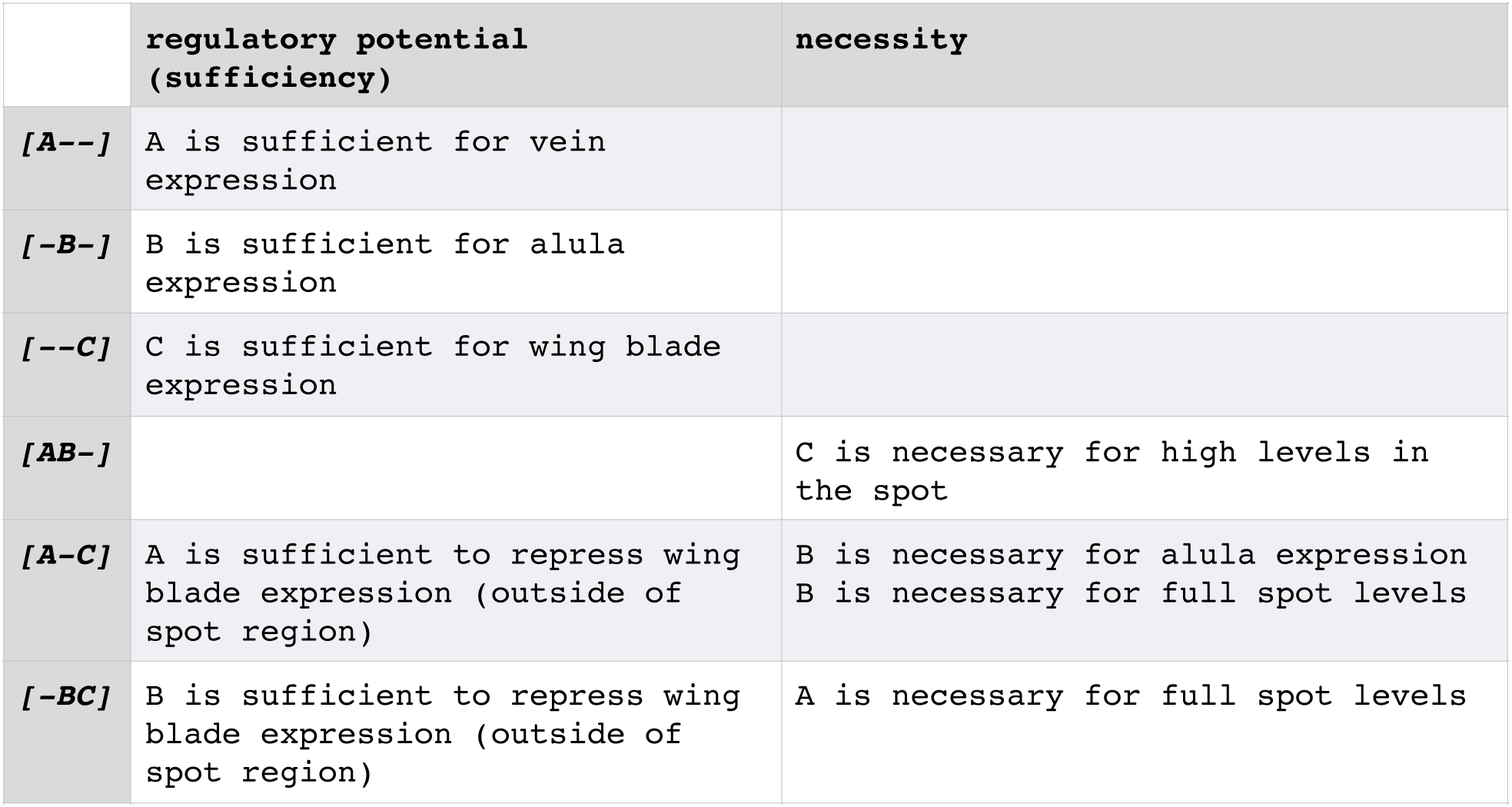
Block contributions. Analysis of necessity and sufficiency of each block.

**Figure 5–figure supplement 2.**
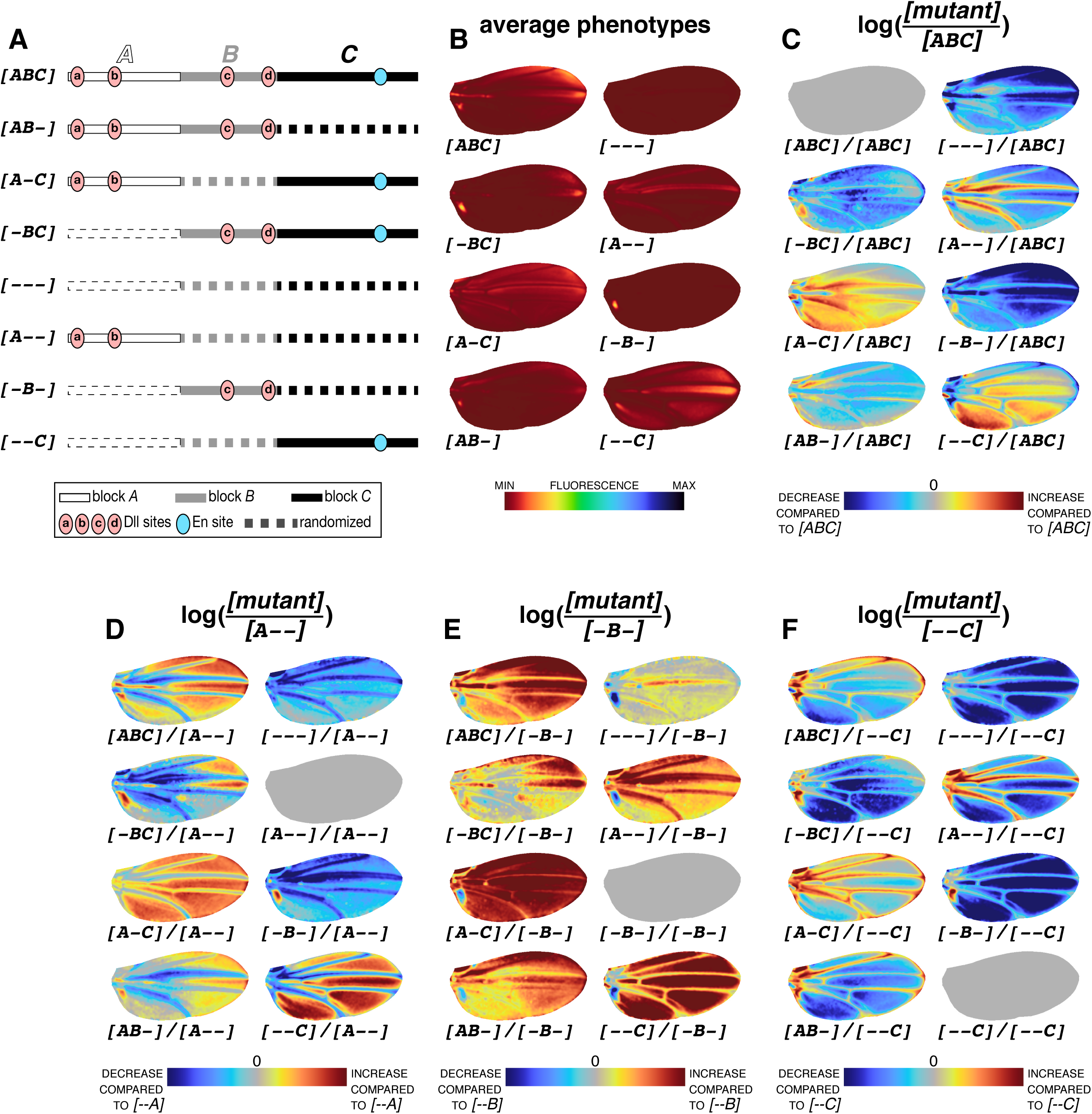
*logRatio* of all block constructs. (**A**) Schematics of block constructs repeated from Figure 5A for legibility. (**B**) Average phenotypes of constructs shown in (**A**), repeated from Figure 5B for legibility. Colormap of average phenotypes normalized for all constructs of the block series, including block permutations of Figure 6B. (**C**) Average phenotypes in (**B**) compared to the average phenotype of the wild type *[ABC]* (*logRatio*). (**D**) Average phenotypes in (**B**) compared to the average phenotype of *[A--]* (*logRatio*). (**E**) Average phenotypes in (**B**) compared to the average phenotype of *[-B-]* (*logRatio*). (**F**) Average phenotypes in (**B**) compared to the average phenotype of *[--C]* (*logRatio*). Colormaps in (**C**)-(**F**) indicate an increase or a decrease of activity compared to the reference (denominator).

**Figure 5–figure supplement 3**. Significance of difference in average expression levels among constructs of the second mutant series (blocks).

## References

(CIE), C.I.d.l.E.c., 2018. Colorimetry, 4th edition ed, CIE Central Bureau, Vienna, Austria.

Abe, N., Dror, I., Yang, L., Slattery, M., Zhou, T., Bussemaker, H.J., Rohs, R., Mann, R.S., 2015. Deconvolving the recognition of DNA shape from sequence. Cell 161, 307–318.

Arnosti, D.N., Barolo, S., Levine, M., Small, S., 1996. The eve stripe 2 enhancer employs multiple modes of transcriptional synergy. Development 122, 205–214.

Arnosti, D.N., Kulkarni, M.M., 2005. Transcriptional enhancers: Intelligent enhanceosomes or flexible billboards? J Cell Biochem 94, 890–898.

Arnoult, L., Su, K.F., Manoel, D., Minervino, C., Magrina, J., Gompel, N., Prud’homme, B., 2013. Emergence and diversification of fly pigmentation through evolution of a gene regulatory module. Science 339, 1423–1426.

Banerji, J., Olson, L., Schaffner, W., 1983. A lymphocyte-specific cellular enhancer is located downstream of the joining region in immunoglobulin heavy chain genes. Cell 33, 729–740.

Barolo, S., 2016. How to tune an enhancer. Proc Natl Acad Sci U S A 113, 6330–6331.

Barolo, S., Castro, B., Posakony, J.W., 2004. New Drosophila transgenic reporters: insulated P-element vectors expressing fast-maturing RFP. Biotechniques 36, 436-440, 442.

Barozzi, I., Simonatto, M., Bonifacio, S., Yang, L., Rohs, R., Ghisletti, S., Natoli, G., 2014. Coregulation of transcription factor binding and nucleosome occupancy through DNA features of mammalian enhancers. Mol Cell 54, 844–857.

Bentovim, L., Harden, T.T., DePace, A.H., 2017. Transcriptional precision and accuracy in development: from measurements to models and mechanisms. Development 144, 3855–3866.

Bertrand, V., Hudson, C., Caillol, D., Popovici, C., Lemaire, P., 2003. Neural tissue in ascidian embryos is induced by FGF9/16/20, acting via a combination of maternal GATA and Ets transcription factors. Cell 115, 615–627.

Carroll, S.B., 1998. From pattern to gene, from gene to pattern. Int J Dev Biol 42, 305–309.

Chiu, T.P., Comoglio, F., Zhou, T., Yang, L., Paro, R., Rohs, R., 2016. DNAshapeR: an R/Bioconductor package for DNA shape prediction and feature encoding. Bioinformatics 32, 1211–1213.

Crocker, J., Abe, N., Rinaldi, L., McGregor, A.P., Frankel, N., Wang, S., Alsawadi, A., Valenti, P., Plaza, S., Payre, F., Mann, R.S., Stern, D.L., 2015. Low affinity binding site clusters confer hox specificity and regulatory robustness. Cell 160, 191–203.

Crocker, J., Stern, D.L., 2017. Functional regulatory evolution outside of the minimal even-skipped stripe 2 enhancer. Development 144, 3095–3101.

Crocker, J., Tsai, A., Stern, D.L., 2017. A Fully Synthetic Transcriptional Platform for a Multicellular Eukaryote. Cell Rep 18, 287–296.

Dufourt, J., Trullo, A., Hunter, J., Fernandez, C., Lazaro, J., Dejean, M., Morales, L., Nait-Amer, S., Schulz, K.N., Harrison, M.M., Favard, C., Radulescu, O., Lagha, M., 2018. Temporal control of gene expression by the pioneer factor Zelda through transient interactions in hubs. Nat Commun 9, 5194.

Erceg, J., Saunders, T.E., Girardot, C., Devos, D.P., Hufnagel, L., Furlong, E.E., 2014. Subtle changes in motif positioning cause tissue-specific effects on robustness of an enhancer’s activity. PLoS Genet 10, e1004060.

Evans, N.C., Swanson, C.I., Barolo, S., 2012. Sparkling insights into enhancer structure, function, and evolution. Curr Top Dev Biol 98, 97–120.

Farley, E.K., Olson, K.M., Zhang, W., Brandt, A.J., Rokhsar, D.S., Levine, M.S., 2015. Suboptimization of developmental enhancers. Science 350, 325–328.

Farley, E.K., Olson, K.M., Zhang, W., Rokhsar, D.S., Levine, M.S., 2016. Syntax compensates for poor binding sites to encode tissue specificity of developmental enhancers. Proc Natl Acad Sci U S A 113, 6508–6513.

Fowlkes, C.C., Hendriks, C.L., Keranen, S.V., Weber, G.H., Rubel, O., Huang, M.Y., Chatoor, S., DePace, A.H., Simirenko, L., Henriquez, C., Beaton, A., Weiszmann, R., Celniker, S., Hamann, B., Knowles, D.W., Biggin, M.D., Eisen, M.B., Malik, J., 2008. A quantitative spatiotemporal atlas of gene expression in the Drosophila blastoderm. Cell 133, 364–374.

Gompel, N., Prud’homme, B., Wittkopp, P.J., Kassner, V.A., Carroll, S.B., 2005. Chance caught on the wing: cis-regulatory evolution and the origin of pigment patterns in Drosophila. Nature 433, 481–487.

Gordan, R., Shen, N., Dror, I., Zhou, T., Horton, J., Rohs, R., Bulyk, M.L., 2013. Genomic regions flanking E-box binding sites influence DNA binding specificity of bHLH transcription factors through DNA shape. Cell Rep 3, 1093–1104.

Gray, S., Levine, M., 1996. Transcriptional repression in development. Curr Opin Cell Biol 8, 358–364.

Gregor, T., Wieschaus, E.F., McGregor, A.P., Bialek, W., Tank, D.W., 2007. Stability and nuclear dynamics of the bicoid morphogen gradient. Cell 130, 141–152.

Groth, A., Fish, M., Nusse, R., Calos, M., 2004. Construction of transgenic Drosophila by using the site-specific integrase from phage phiC31. Genetics 166, 1775–1782.

Hare, E.E., Peterson, B.K., Eisen, M.B., 2008. A careful look at binding site reorganization in the even-skipped enhancers of Drosophila and sepsids. PLoS Genet 4, e1000268.

Hizver, J., Rozenberg, H., Frolow, F., Rabinovich, D., Shakked, Z., 2001. DNA bending by an adenine--thymine tract and its role in gene regulation. Proc Natl Acad Sci U S A 98, 8490–8495.

Hutchinson, M.F., 1995. Interpolating mean rainfall using thin plate smoothing splines. International Journal of Geographical Information Systems 9, 385–403.

Khoueiry, P., Rothbacher, U., Ohtsuka, Y., Daian, F., Frangulian, E., Roure, A., Dubchak, I., Lemaire, P., 2010. A cis-regulatory signature in ascidians and flies, independent of transcription factor binding sites. Curr Biol 20, 792–802.

King, D.M., Hong, C.K.Y., Shepherdson, J.L., Granas, D.M., Maricque, B.B., Cohen, B.A., 2020. Synthetic and genomic regulatory elements reveal aspects of cis-regulatory grammar in mouse embryonic stem cells. Elife 9.

Kircher, M., Xiong, C., Martin, B., Schubach, M., Inoue, F., Bell, R.J.A., Costello, J.F., Shendure, J., Ahituv, N., 2019. Saturation mutagenesis of twenty disease-associated regulatory elements at single base-pair resolution. Nat Commun 10, 3583.

Knowles, D.W., 2012. Three-dimensional morphology and gene expression mapping for the Drosophila blastoderm. Cold Spring Harb Protoc 2012, 150–161.

Kulkarni, M.M., Arnosti, D.N., 2003. Information display by transcriptional enhancers. Development 130, 6569–6575.

Kwasnieski, J.C., Mogno, I., Myers, C.A., Corbo, J.C., Cohen, B.A., 2012. Complex effects of nucleotide variants in a mammalian cis-regulatory element. Proc Natl Acad Sci U S A 109, 19498–19503.

Le Poul, Y., Bahry, E., Gompel, N., Preibisch, S., in preparation. Active appearance model for automated landmark and sliding landmark positioning.

Levine, M., 2010. Transcriptional enhancers in animal development and evolution. Curr Biol 20, R754–763.

Levo, M., Segal, E., 2014. In pursuit of design principles of regulatory sequences. Nat Rev Genet 15, 453–468.

Lim, F.L., Hayes, A., West, A.G., Pic-Taylor, A., Darieva, Z., Morgan, B.A., Oliver, S.G., Sharrocks, A.D., 2003. Mcm1p-induced DNA bending regulates the formation of ternary transcription factor complexes. Mol Cell Biol 23, 450–461.

Long, H.K., Prescott, S.L., Wysocka, J., 2016. Ever-Changing Landscapes: Transcriptional Enhancers in Development and Evolution. Cell 167, 1170–1187.

Ludwig, M.Z., Bergman, C., Patel, N.H., Kreitman, M., 2000. Evidence for stabilizing selection in a eukaryotic enhancer element. Nature 403, 564–567.

Ludwig, M.Z., Manu Kittler, R., White, K.P., Kreitman, M., 2011. Consequences of eukaryotic enhancer architecture for gene expression dynamics, development, and fitness. PLoS Genet 7, e1002364.

Ludwig, M.Z., Palsson, A., Alekseeva, E., Bergman, C.M., Nathan, J., Kreitman, M., 2005. Functional evolution of a cis-regulatory module. PLoS Biol 3, e93.

Martinez-Abadias, N., Mateu Estivill, R., Sastre Tomas, J., Motch Perrine, S., Yoon, M., Robert-Moreno, A., Swoger, J., Russo, L., Kawasaki, K., Richtsmeier, J., Sharpe, J., 2018. Quantification of gene expression patterns to reveal the origins of abnormal morphogenesis. Elife 7.

Martinez-Abadias, N., Mateu, R., Niksic, M., Russo, L., Sharpe, J., 2016. Geometric Morphometrics on Gene Expression Patterns Within Phenotypes: A Case Example on Limb Development. Syst Biol 65, 194–211.

Morgunova, E., Taipale, J., 2017. Structural perspective of cooperative transcription factor binding. Current opinion in structural biology 47, 1–8.

Neidle, S., 2010. Principles of Nucleic Acid Structure. Academic Press.

Nelson, H.C., Finch, J.T., Luisi, B.F., Klug, A., 1987. The structure of an oligo(dA).oligo(dT) tract and its biological implications. Nature 330, 221–226.

Nitzan, M., Karaiskos, N., Friedman, N., Rajewsky, N., 2019. Gene expression cartography. Nature 576, 132–137.

Panne, D., Maniatis, T., Harrison, S.C., 2007. An atomic model of the interferon-beta enhanceosome. Cell 129, 1111–1123.

Park, J., Estrada, J., Johnson, G., Vincent, B.J., Ricci-Tam, C., Bragdon, M.D., Shulgina, Y., Cha, A., Wunderlich, Z., Gunawardena, J., DePace, A.H., 2019. Dissecting the sharp response of a canonical developmental enhancer reveals multiple sources of cooperativity. Elife 8.

Peter, I.S., Davidson, E.H., 2015. Genomic Control Process: Development and Evolution, 1st ed. Academic Press, San Diego, United States.

Robinson, M.D., McCarthy, D.J., Smyth, G.K., 2010. edgeR: a Bioconductor package for differential expression analysis of digital gene expression data. Bioinformatics 26, 139–140.

Rohs, R., West, S.M., Sosinsky, A., Liu, P., Mann, R.S., Honig, B., 2009. The role of DNA shape in protein-DNA recognition. Nature 461, 1248–1253.

Ronchi, E., Treisman, J., Dostatni, N., Struhl, G., Desplan, C., 1993. Down-regulation of the Drosophila morphogen bicoid by the torso receptor-mediated signal transduction cascade. Cell 74, 347–355.

Shlyueva, D., Stampfel, G., Stark, A., 2014. Transcriptional enhancers: from properties to genome-wide predictions. Nat Rev Genet 15, 272–286.

Slattery, M., Zhou, T., Yang, L., Dantas Machado, A.C., Gordan, R., Rohs, R., 2014. Absence of a simple code: how transcription factors read the genome. Trends Biochem Sci 39, 381–399.

Spitz, F., Furlong, E.E., 2012. Transcription factors: from enhancer binding to developmental control. Nat Rev Genet 13, 613–626.

Stewart, A.J., Hannenhalli, S., Plotkin, J.B., 2012. Why transcription factor binding sites are ten nucleotides long. Genetics 192, 973–985.

Suter, B., Schnappauf, G., Thoma, F., 2000. Poly(dA.dT) sequences exist as rigid DNA structures in nucleosome-free yeast promoters in vivo. Nucleic Acids Res 28, 4083–4089.

Swanson, C.I., Evans, N.C., Barolo, S., 2010. Structural rules and complex regulatory circuitry constrain expression of a Notch- and EGFR-regulated eye enhancer. Dev Cell 18, 359–370.

Swanson, C.I., Schwimmer, D.B., Barolo, S., 2011. Rapid evolutionary rewiring of a structurally constrained eye enhancer. Curr Biol 21, 1186–1196.

Thanos, D., Maniatis, T., 1995. Virus induction of human IFN beta gene expression requires the assembly of an enhanceosome. Cell 83, 1091–1100.

Venken, K.J., He, Y., Hoskins, R.A., Bellen, H.J., 2006. P[acman]: a BAC transgenic platform for targeted insertion of large DNA fragments in D. melanogaster. Science 314, 1747–1751.

Vincent, B.J., Estrada, J., DePace, A.H., 2016. The appeasement of Doug: a synthetic approach to enhancer biology. Integr Biol (Camb) 8, 475–484.

Weingarten-Gabbay, S., Segal, E., 2014. The grammar of transcriptional regulation. Hum Genet 133, 701–711.

Wittkopp, P.J., Carroll, S.B., Kopp, A., 2003. Evolution in black and white: genetic control of pigment patterns in Drosophila. Trends Genet 19, 495–504.

Xin, Y., Le Poul, Y., Ling, L., Museridze, M., Mühling, B., Jaenichen, R., Osipova, E., Gompel, N., 2020. An ancestral and a derived transcriptional enhancers sharing regulatory sequence and pleiotropic control of chromatin accessibility. submitted.

Yanez-Cuna, J.O., Kvon, E.Z., Stark, A., 2013. Deciphering the transcriptional cis-regulatory code. Trends Genet 29, 11–22.

Yella, V.R., Bhimsaria, D., Ghoshdastidar, D., Rodriguez-Martinez, J.A., Ansari, A.Z., Bansal, M., 2018. Flexibility and structure of flanking DNA impact transcription factor affinity for its core motif. Nucleic Acids Res 46, 11883–11897.

Zhou, T., Yang, L., Lu, Y., Dror, I., Dantas Machado, A.C., Ghane, T., Di Felice, R., Rohs, R., 2013. DNAshape: a method for the high-throughput prediction of DNA structural features on a genomic scale. Nucleic Acids Res 41, W56–62.

Zhu, L.J., Christensen, R.G., Kazemian, M., Hull, C.J., Enuameh, M.S., Basciotta, M.D., Brasefield, J.A., Zhu, C., Asriyan, Y., Lapointe, D.S., Sinha, S., Wolfe, S.A., Brodsky, M.H., 2011. FlyFactorSurvey: a database of Drosophila transcription factor binding specificities determined using the bacterial one-hybrid system. Nucleic Acids Res 39, D111–117.

