## Supplemental Data 1 for "Deciphering the regulatory logic of a *Drosophila* enhancer through systematic sequence mutagenesis and quantitative image analysis"

**Le Poul, Xin et al. – Figure 1–figure supplement 1**. Sequences of *spot^196^* enhancer variants.

- wild type *[+]* or *[ABC]*

>*spot^196 [+]^*

TCTAATTATTCCGTTTAAGGACGCAATTTTCTGAGCTAAAACTCGCTTATGGAGAGATCTAAATTTCCCCGCTTTTGGCTTGAATAAATTAATCGAATTCCCCGCTGGCTATTAAAACACACAAAAGGCGCTCTCGTCTGTTTCAATGTAAATTGCAAATTGCTCAATCCGCCTAATTGATGTGCGCCCATGCAAT

- single mutants *[0]* to *[16]*

>*spot^196 [0]^*

AAAAAAAAAAAAAAAAAAGGACGCAATTTTCTGAGCTAAAACTCGCTTATGGAGAGATCTAAATTTCCCCGCTTTTGGCTTGAATAAATTAATCGAATTCCCCGCTGGCTATTAAAACACACAAAAGGCGCTCTCGTCTGTTTCAATGTAAATTGCAAATTGCTCAATCCGCCTAATTGATGTGCGCCCATGCAAT

>*spot^196 [1]^*

TCTAATTATTCCGTTTAAAAAAAAAAAATTCTGAGCTAAAACTCGCTTATGGAGAGATCTAAATTTCCCCGCTTTTGGCTTGAATAAATTAATCGAATTCCCCGCTGGCTATTAAAACACACAAAAGGCGCTCTCGTCTGTTTCAATGTAAATTGCAAATTGCTCAATCCGCCTAATTGATGTGCGCCCATGCAAT

>*spot^196 [2]^*

TCTAATTATTCCGTTTAAGGACGCAATTAAAAAAAAAAAAACTCGCTTATGGAGAGATCTAAATTTCCCCGCTTTTGGCTTGAATAAATTAATCGAATTCCCCGCTGGCTATTAAAACACACAAAAGGCGCTCTCGTCTGTTTCAATGTAAATTGCAAATTGCTCAATCCGCCTAATTGATGTGCGCCCATGCAAT

>*spot^196 [3]^*

TCTAATTATTCCGTTTAAGGACGCAATTTTCTGAGCTAAAAAAAAAAAATGGAGAGATCTAAATTTCCCCGCTTTTGGCTTGAATAAATTAATCGAATTCCCCGCTGGCTATTAAAACACACAAAAGGCGCTCTCGTCTGTTTCAATGTAAATTGCAAATTGCTCAATCCGCCTAATTGATGTGCGCCCATGCAAT

>*spot^196 [4]^*

TCTAATTATTCCGTTTAAGGACGCAATTTTCTGAGCTAAAACTCGCTTAAAAAAAAAAATAAATTTCCCCGCTTTTGGCTTGAATAAATTAATCGAATTCCCCGCTGGCTATTAAAACACACAAAAGGCGCTCTCGTCTGTTTCAATGTAAATTGCAAATTGCTCAATCCGCCTAATTGATGTGCGCCCATGCAAT

>*spot^196 [5]^*

TCTAATTATTCCGTTTAAGGACGCAATTTTCTGAGCTAAAACTCGCTTATGGAGAGATCAAAAAAAAAAAGCTTTTGGCTTGAATAAATTAATCGAATTCCCCGCTGGCTATTAAAACACACAAAAGGCGCTCTCGTCTGTTTCAATGTAAATTGCAAATTGCTCAATCCGCCTAATTGATGTGCGCCCATGCAAT

>*spot^196 [6]^*

TCTAATTATTCCGTTTAAGGACGCAATTTTCTGAGCTAAAACTCGCTTATGGAGAGATCTAAATTTCCCCAAAAAAAAAAAAAATAAATTAATCGAATTCCCCGCTGGCTATTAAAACACACAAAAGGCGCTCTCGTCTGTTTCAATGTAAATTGCAAATTGCTCAATCCGCCTAATTGATGTGCGCCCATGCAAT

>*spot^196 [7]^*

TCTAATTATTCCGTTTAAGGACGCAATTTTCTGAGCTAAAACTCGCTTATGGAGAGATCTAAATTTCCCCGCTTTTGGCTTGAAAAAAAAAAAAGAATTCCCCGCTGGCTATTAAAACACACAAAAGGCGCTCTCGTCTGTTTCAATGTAAATTGCAAATTGCTCAATCCGCCTAATTGATGTGCGCCCATGCAAT

>*spot^196 [8]^*

TCTAATTATTCCGTTTAAGGACGCAATTTTCTGAGCTAAAACTCGCTTATGGAGAGATCTAAATTTCCCCGCTTTTGGCTTGAATAAATTAATCAAAAAAAAAAAAGGCTATTAAAACACACAAAAGGCGCTCTCGTCTGTTTCAATGTAAATTGCAAATTGCTCAATCCGCCTAATTGATGTGCGCCCATGCAAT

>*spot^196 [9]^*

TCTAATTATTCCGTTTAAGGACGCAATTTTCTGAGCTAAAACTCGCTTATGGAGAGATCTAAATTTCCCCGCTTTTGGCTTGAATAAATTAATCGAATTCCCCGCTAAAAAAAAAAACACACAAAAGGCGCTCTCGTCTGTTTCAATGTAAATTGCAAATTGCTCAATCCGCCTAATTGATGTGCGCCCATGCAAT

>*spot^196 [10]^*

TCTAATTATTCCGTTTAAGGACGCAATTTTCTGAGCTAAAACTCGCTTATGGAGAGATCTAAATTTCCCCGCTTTTGGCTTGAATAAATTAATCGAATTCCCCGCTGGCTATTAAAAAAAAAAAAAGGCGCTCTCGTCTGTTTCAATGTAAATTGCAAATTGCTCAATCCGCCTAATTGATGTGCGCCCATGCAAT

>*spot^196 [11]^*

TCTAATTATTCCGTTTAAGGACGCAATTTTCTGAGCTAAAACTCGCTTATGGAGAGATCTAAATTTCCCCGCTTTTGGCTTGAATAAATTAATCGAATTCCCCGCTGGCTATTAAAACACACAAAAAAAAAAAAAATCTGTTTCAATGTAAATTGCAAATTGCTCAATCCGCCTAATTGATGTGCGCCCATGCAAT

>*spot^196 [12]^*

TCTAATTATTCCGTTTAAGGACGCAATTTTCTGAGCTAAAACTCGCTTATGGAGAGATCTAAATTTCCCCGCTTTTGGCTTGAATAAATTAATCGAATTCCCCGCTGGCTATTAAAACACACAAAAGGCGCTCTCGAAAAAAAAAATGTAAATTGCAAATTGCTCAATCCGCCTAATTGATGTGCGCCCATGCAAT

>*spot^196 [13]^*

TCTAATTATTCCGTTTAAGGACGCAATTTTCTGAGCTAAAACTCGCTTATGGAGAGATCTAAATTTCCCCGCTTTTGGCTTGAATAAATTAATCGAATTCCCCGCTGGCTATTAAAACACACAAAAGGCGCTCTCGTCTGTTTCAAAAAAAAAAAAAAATTGCTCAATCCGCCTAATTGATGTGCGCCCATGCAAT

>*spot^196 [14]^*

TCTAATTATTCCGTTTAAGGACGCAATTTTCTGAGCTAAAACTCGCTTATGGAGAGATCTAAATTTCCCCGCTTTTGGCTTGAATAAATTAATCGAATTCCCCGCTGGCTATTAAAACACACAAAAGGCGCTCTCGTCTGTTTCAATGTAAATTGCAAAAAAAAAAAACCGCCTAATTGATGTGCGCCCATGCAAT

>*spot^196 [15]^*

TCTAATTATTCCGTTTAAGGACGCAATTTTCTGAGCTAAAACTCGCTTATGGAGAGATCTAAATTTCCCCGCTTTTGGCTTGAATAAATTAATCGAATTCCCCGCTGGCTATTAAAACACACAAAAGGCGCTCTCGTCTGTTTCAATGTAAATTGCAAATTGCTCAATAAAAAAAAAAAATGTGCGCCCATGCAAT

>*spot^196 [16]^*

TCTAATTATTCCGTTTAAGGACGCAATTTTCTGAGCTAAAACTCGCTTATGGAGAGATCTAAATTTCCCCGCTTTTGGCTTGAATAAATTAATCGAATTCCCCGCTGGCTATTAAAACACACAAAAGGCGCTCTCGTCTGTTTCAATGTAAATTGCAAATTGCTCAATCCGCCTAATTGAAAAAAAAAAAAAAAAA

- Permutations of blocks

> *spot^196 [AC^**^B]^*

TCTAATTATTCCGTTTAAGGACGCAATTTTCTGAGCTAAAACTCGCTTATGGAGAGATCTAAACACACAAAAGGCGCTCTCGTCTGTTTCAATGTAAATTGCAAATTGCTCAATCCGCCTAATTGATGTGCGCCCATGCAATTTTCCCCGCTTTTGGCTTGAATAAATTAATCGAATTCCCCGCTGGCTATTAAAA

>*spot^196 [BAC]^*

TTTCCCCGCTTTTGGCTTGAATAAATTAATCGAATTCCCCGCTGGCTATTAAAATCTAATTATTCCGTTTAAGGACGCAATTTTCTGAGCTAAAACTCGCTTATGGAGAGATCTAAACACACAAAAGGCGCTCTCGTCTGTTTCAATGTAAATTGCAAATTGCTCAATCCGCCTAATTGATGTGCGCCCATGCAAT

>*spot^196 [BCA]^*

TTTCCCCGCTTTTGGCTTGAATAAATTAATCGAATTCCCCGCTGGCTATTAAAACACACAAAAGGCGCTCTCGTCTGTTTCAATGTAAATTGCAAATTGCTCAATCCGCCTAATTGATGTGCGCCCATGCAATTCTAATTATTCCGTTTAAGGACGCAATTTTCTGAGCTAAAACTCGCTTATGGAGAGATCTAAA

>*spot^196 [CBA]^*

CACACAAAAGGCGCTCTCGTCTGTTTCAATGTAAATTGCAAATTGCTCAATCCGCCTAATTGATGTGCGCCCATGCAATTTTCCCCGCTTTTGGCTTGAATAAATTAATCGAATTCCCCGCTGGCTATTAAAATCTAATTATTCCGTTTAAGGACGCAATTTTCTGAGCTAAAACTCGCTTATGGAGAGATCTAAA

- Randomized blocks

>*spot^196 [A--]^*

TCTAATTATTCCGTTTAAGGACGCAATTTTCTGAGCTAAAACTCGCTTATGGAGAGATCTAAATCCGAATTTTTTCTTGTCCGACTAGAAACGACTAATTTAGCCGTACCACATGTTGTCGACTCAGAAACATTATTCCCATTTACGCGTAAGCAAAAAATGCGTCCTTATCGAACTTACACTCGCCTGCGTTGGT

>*spot^196 [-B-]^*

ATAATATTGCATCTCATTGTGGTGCTAGATAATCATCTAGGCTAAATCCAAAACTGTTGCATGTTTCCCCGCTTTTGGCTTGAATAAATTAATCGAATTCCCCGCTGGCTATTAAAAGTCGACTCAGAAACATTATTCCCATTTACGCGTAAGCAAAAAATGCGTCCTTATCGAACTTACACTCGCCTGCGTTGGT

>*spot^196 [--C]^*

ATAATATTGCATCTCATTGTGGTGCTAGATAATCATCTAGGCTAAATCCAAAACTGTTGCATGTCCGAATTTTTTCTTGTCCGACTAGAAACGACTAATTTAGCCGTACCACATGTTCACACAAAAGGCGCTCTCGTCTGTTTCAATGTAAATTGCAAATTGCTCAATCCGCCTAATTGATGTGCGCCCATGCAAT

>*spot^196 [AB-]^*

TCTAATTATTCCGTTTAAGGACGCAATTTTCTGAGCTAAAACTCGCTTATGGAGAGATCTAAATTTCCCCGCTTTTGGCTTGAATAAATTAATCGAATTCCCCGCTGGCTATTAAAAGTCGACTCAGAAACATTATTCCCATTTACGCGTAAGCAAAAAATGCGTCCTTATCGAACTTACACTCGCCTGCGTTGGT

>*spot^196 [A-C]^*

TCTAATTATTCCGTTTAAGGACGCAATTTTCTGAGCTAAAACTCGCTTATGGAGAGATCTAAATCCGAATTTTTTCTTGTCCGACTAGAAACGACTAATTTAGCCGTACCACATGTTCACACAAAAGGCGCTCTCGTCTGTTTCAATGTAAATTGCAAATTGCTCAATCCGCCTAATTGATGTGCGCCCATGCAAT

>*spot^196 [-BC]^*

ATAATATTGCATCTCATTGTGGTGCTAGATAATCATCTAGGCTAAATCCAAAACTGTTGCATGTTTCCCCGCTTTTGGCTTGAATAAATTAATCGAATTCCCCGCTGGCTATTAAAACACACAAAAGGCGCTCTCGTCTGTTTCAATGTAAATTGCAAATTGCTCAATCCGCCTAATTGATGTGCGCCCATGCAAT

>*spot^196 [---]^*

ATAATATTGCATCTCATTGTGGTGCTAGATAATCATCTAGGCTAAATCCAAAACTGTTGCATGTCCGAATTTTTTCTTGTCCGACTAGAAACGACTAATTTAGCCGTACCACATGTTGTCGACTCAGAAACATTATTCCCATTTACGCGTAAGCAAAAAATGCGTCCTTATCGAACTTACACTCGCCTGCGTTGGT
